# Computational design of HLA class I superbinders for broad T cell immunogenicity

**DOI:** 10.1101/2025.07.10.664084

**Authors:** Elinor Peer, Liel Cohen-Lavi, Alessandro Sette, John Sidney, Tomer Hertz

## Abstract

Human leukocyte antigen (HLA) class I molecules are highly polymorphic, restricting peptide binding to narrow sequence subsets. Designing peptides that bind multiple HLA supertypes— termed superbinders—offers a promising strategy for broad-spectrum T-cell vaccines and immunotherapies. Here we present superHLA, a computational framework that combines Markov Chain Monte Carlo optimization with state-of-the-art MHC binding predictors to design synthetic 9-mer peptides with broad HLA-binding profiles. Using superHLA, we generated over 190,000 candidate superbinders predicted to bind 8–12 HLA class I alleles across distinct supertypes. A multi-tier filtering pipeline—incorporating sequence clustering, synthesis feasibility, cross-predictor validation, and self-peptidome exclusion—yielded a final panel of 100 peptides for experimental testing. Of these, 21 bound ≥4 supertypes in vitro, including one that bound 9. Superbinders displayed distinct anchor residue preferences and showed minimal similarity to human peptides. These results suggest that HLA superbinders are more abundant than previously recognized and can be rationally designed at scale. This approach supports development of pan-HLA immunogens with broad population coverage and potential applications in vaccine design, neoantigen discovery, and immunotherapy.

## Introduction

Most adaptive immune system effector functions are initiated by the Major Histocompatibility Complex (MHC) pathway ^1^, which presents self-peptides and pathogenic peptides from within the cell on the cell surface for surveillance by T cells^2^. The interactions between MHC molecules and peptides and subsequent recognition of these MHC-peptide complexes by T cells control the magnitude and effectiveness of the T-cell-mediated adaptive immune response^3^. In humans, MHC alleles, also known as human leukocyte antigen (HLA), have evolved to optimize the fitness of human populations in the face of a wide range of pathogen species and the broad spectrum of genetic variations that some of these pathogens exhibit.

The HLA gene complex is among the most genetically diverse regions in the human genome, with more than 22,000 HLA class I alleles identified to date^4^. This diversity is driven and maintained by both heterozygote advantage and frequency-dependent selection, which enables low-frequency alleles to gain a fitness advantage as the spectrum of circulating pathogens changes over time^5,6^. HLA molecules are highly selective and only bind to specific peptides. Each molecule is estimated to bind ∼1% of all existing peptides^7^. Polymorphisms in HLA genes have been shown to contribute to individual fitness under natural and sexual selection^1,8^. Despite their broad diversity, HLA alleles can be grouped into Supertypes - HLA alleles that have similar peptide binding specificities^9–11^.

The peptide repertoire presented by HLA proteins largely depends on the structural features of the binding groove of each particular HLA allelic variant. The amino acid structure of each binding pocket thus determines the peptide side chain specificity. In the case of HLA class I, the main binding energy is provided by the interaction of residues in position 2 and the C-terminus (position 9) of the peptide with the B and F binding pockets of the HLA molecule, respectively^12,9,13^. While a typical antigen encodes hundreds of 9-mer peptides, only a small fraction of these will bind HLA alleles^1,4^. These immunodominant regions have been previously shown to be enriched in evolutionarily conserved regions of viral and self-antigens^14^. Furthermore, only a fraction of these peptide-HLA complexes will subsequently induce a functional T-cell response^15^.

While each HLA binds to a specific set of peptides, previous work has reported the existence of epitope hotspots - which are defined as public immunodominant regions that may contain several epitopes that are presented by different HLA alleles and are targeted by many individuals ^16,17^. These regions typically encompass 12-20 amino acids, including many overlapping HLA binding sites. While the majority of peptides that have been experimentally shown to bind HLA class I alleles are reported to bind a single HLA allele or multiple alleles from the same supertype, some peptides have been reported to bind alleles from several HLA supertypes, defined here as *superbinders*^18,16^. Currently, only 56 of the ∼38,000 peptides in the Immune epitope database (IEDB) that were experimentally determined to bind HLA are reported to bind more than 5 HLA class I alleles from different supertypes^19^. Therefore, while the existence of superbinders has been demonstrated experimentally, it is unclear how common these peptides are, and how additional superbinders can be systematically discovered.

The development of machine learning-based models and a growing amount of experimental HLA peptide binding data has enabled the development of state-of-the-art HLA binding predictors that predict which peptides can bind to a specific HLA allele, including alleles with little or no available experimental binding data^3,20–23^. Building on these prediction models, we developed *superHLA* - a novel computational approach for identifying HLA superbinders - 9-mer peptides that bind to a broad set of HLA class I alleles from different HLA supertypes. We used our method to generate a large number of putative HLA superbinders *in-silico*, and then validated 100 of these predicted superbinders using biochemical HLA binding assays. We were able to identify 10 novel peptides that bound more than 5 HLA alleles from different supertypes. Our results suggest that HLA superbinders are more prevalent the previously described and can be readily identified using the *superHLA* algorithm.

## Results

### The super-HLA algorithm

To identify novel HLA class I superbinders we developed *superHLA* - an in-silico method for generating a putative list of superbinders - peptides that are predicted to bind to multiple HLA alleles from different HLA class I supertypes (**Figure 1**). To generate putative superbinders, we used a Markov Chain Monte Carlo (MCMC) algorithm to sample the 9-mer peptide space. ^24^ Overall, there are 20^9^unique 9-mer peptides, from which only a small subset binds HLA class I alleles, and an even smaller subset binds multiple HLA class I alleles. To explore this vast space, we used a random walk, in which each step attempts to maximize a scoring function based on the predicted binding affinity of the given peptide. At every simulation step we introduce a single point mutation into the current peptide sequence, aimed at increasing its binding affinity to 12 HLA class I alleles, each from a different supertype **(Figure 2A)**. Each supertype was represented using its most frequent allele (**Table 1**)^25^. The percentile rank scores of the original and mutated peptide are predicted for all 12 HLA alleles^26^. These scores are then aggregated into a global score estimating the predicted change in HLA binding induced by the suggested mutation in the current step. The step is then accepted or rejected by drawing a single value from the uniform distribution in the interval (0, 1) using the acceptance probability **(Algorithm 1 2.2.5)** as a threshold value. Any value below or equal to the threshold denotes acceptance of the mutation, while any number above the threshold denotes rejection of the mutation (**Figure 2B**). The acceptance probability function was selected to assign a high probability to changes that increase binding affinity. In other words, high acceptance probabilities are assigned to moves in which the mutated peptide increases the number of HLA supertypes it binds compared to the previous peptide (**Figure 2B** and **Figure 3**). However, it also allows for slight decreases in binding affinity, allowing the procedure to escape local maxima (**Figure 3**). The number of putative superbinder peptides generated depends on the number of iterations performed by the algorithm.

**Figure 1.**
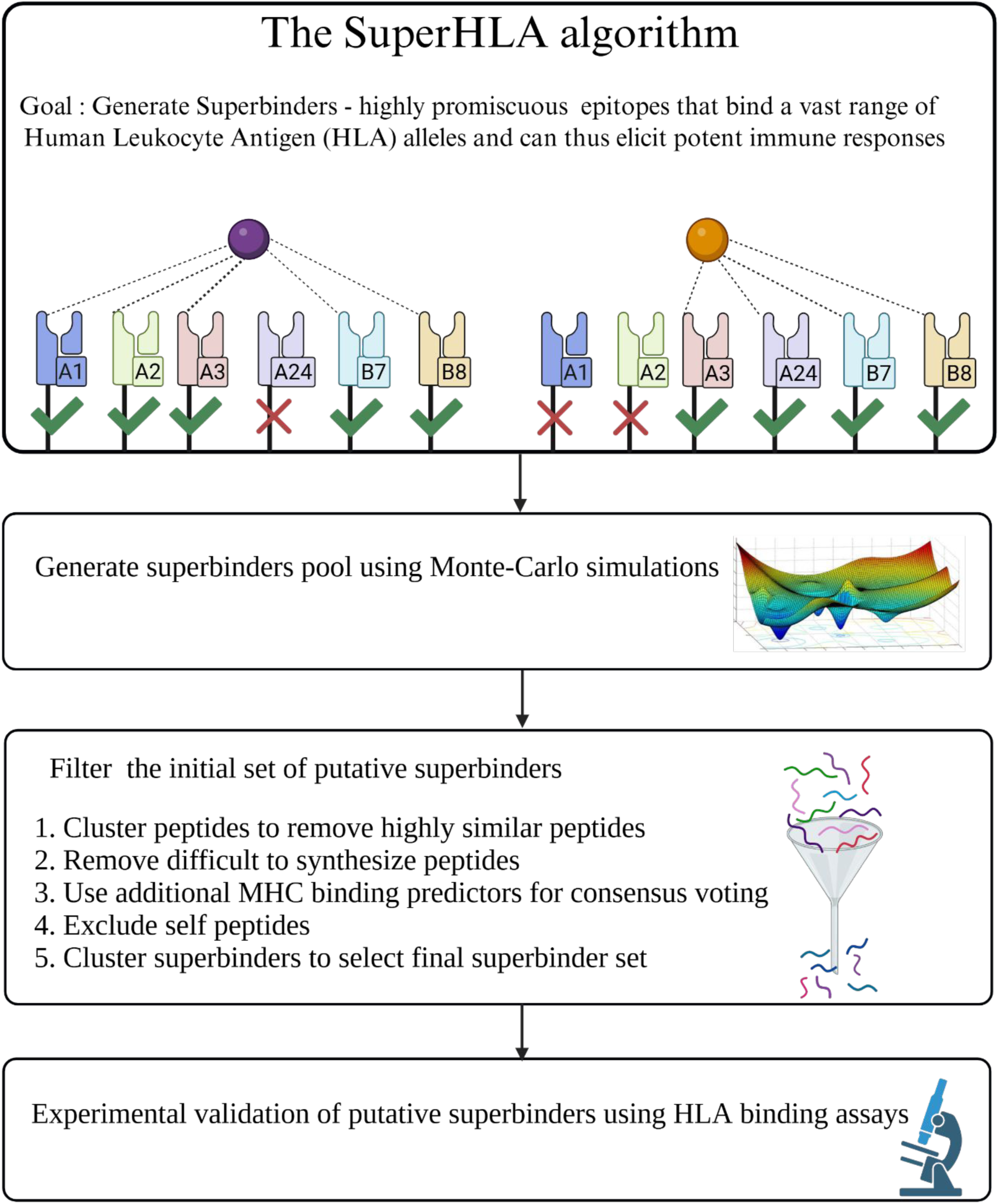
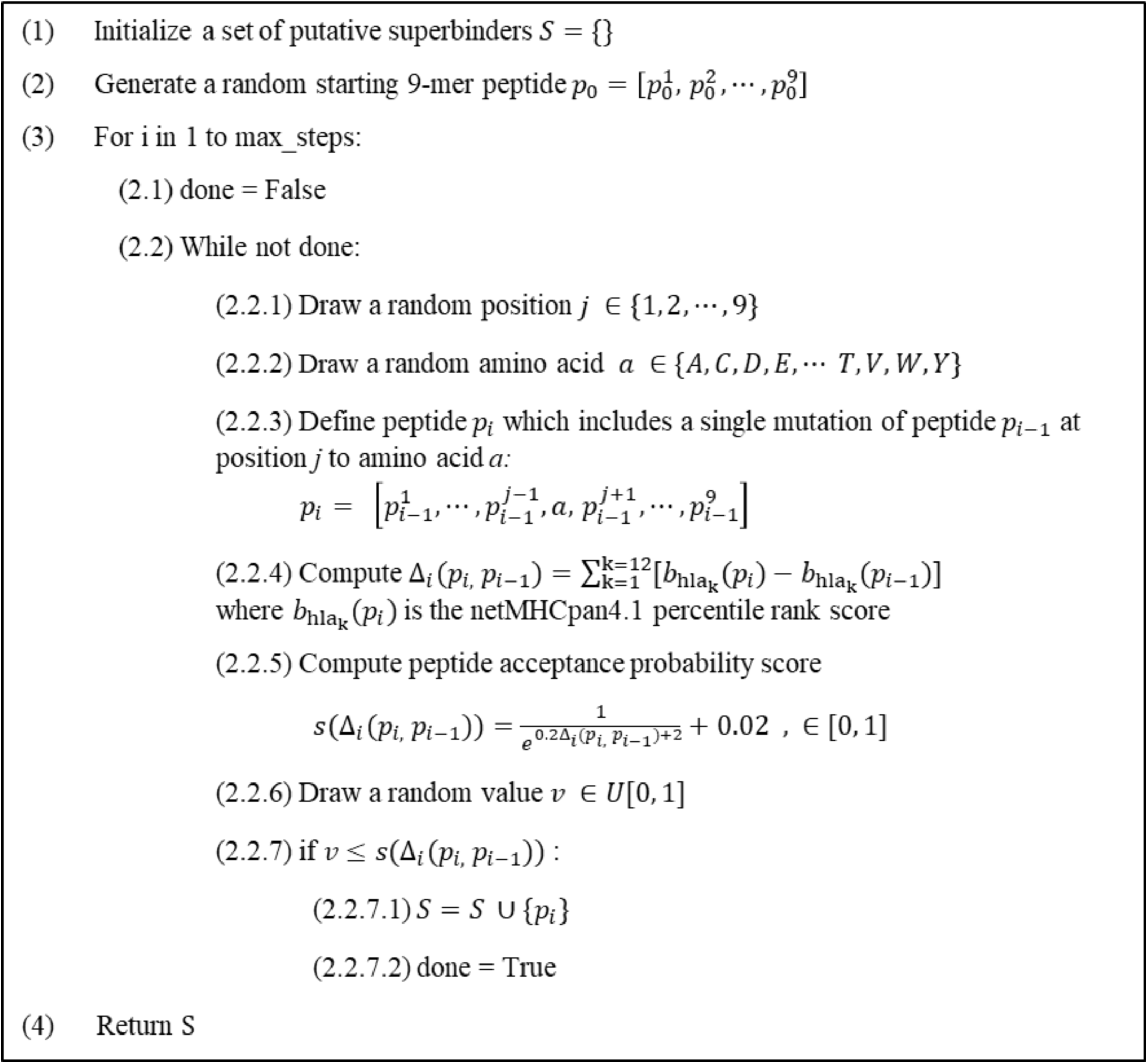
The *SuperHLA* Algorithm.

**Figure 2.**
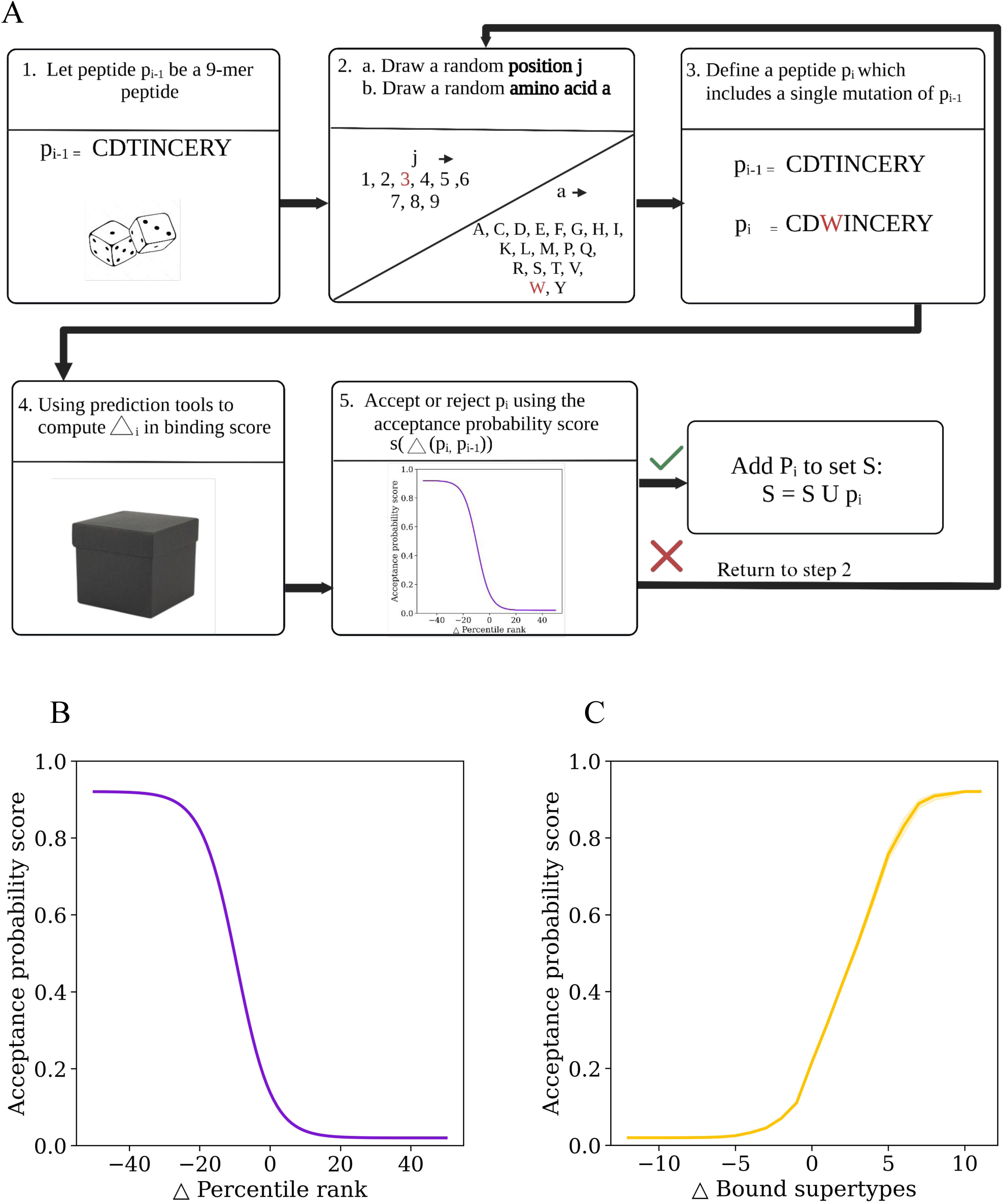
A single simulation step of the super-HLA algorithm. **(A) Workflow of a single simulation step.** Sampling the distribution space while using acceptance probability score enables the *super-HLA* algorithm to also accept steps in which the overall binding affinity is reduced, allowing it to escape local maxima. **(B)** A**cceptance probability score function of the super-HLA algorithm.** The algorithm uses an acceptance probability score function to assign a probability to each simulation step, which assigns higher probabilities to steps that improve the overall binding affinity across the 12 HLA class I supertype representatives. The acceptance probability score s is a function of the Δ in percentile rank - defined as the difference in the HLA binding percentile rank of the peptide and its single-point mutant across all 12 representative HLA alleles. **(C) Acceptance probability as a function of the difference in number of bound supertypes**. As expected, as the number of supertypes a peptide binds increases, the likelihood of accepting the peptide also increases.

**Figure 3.**
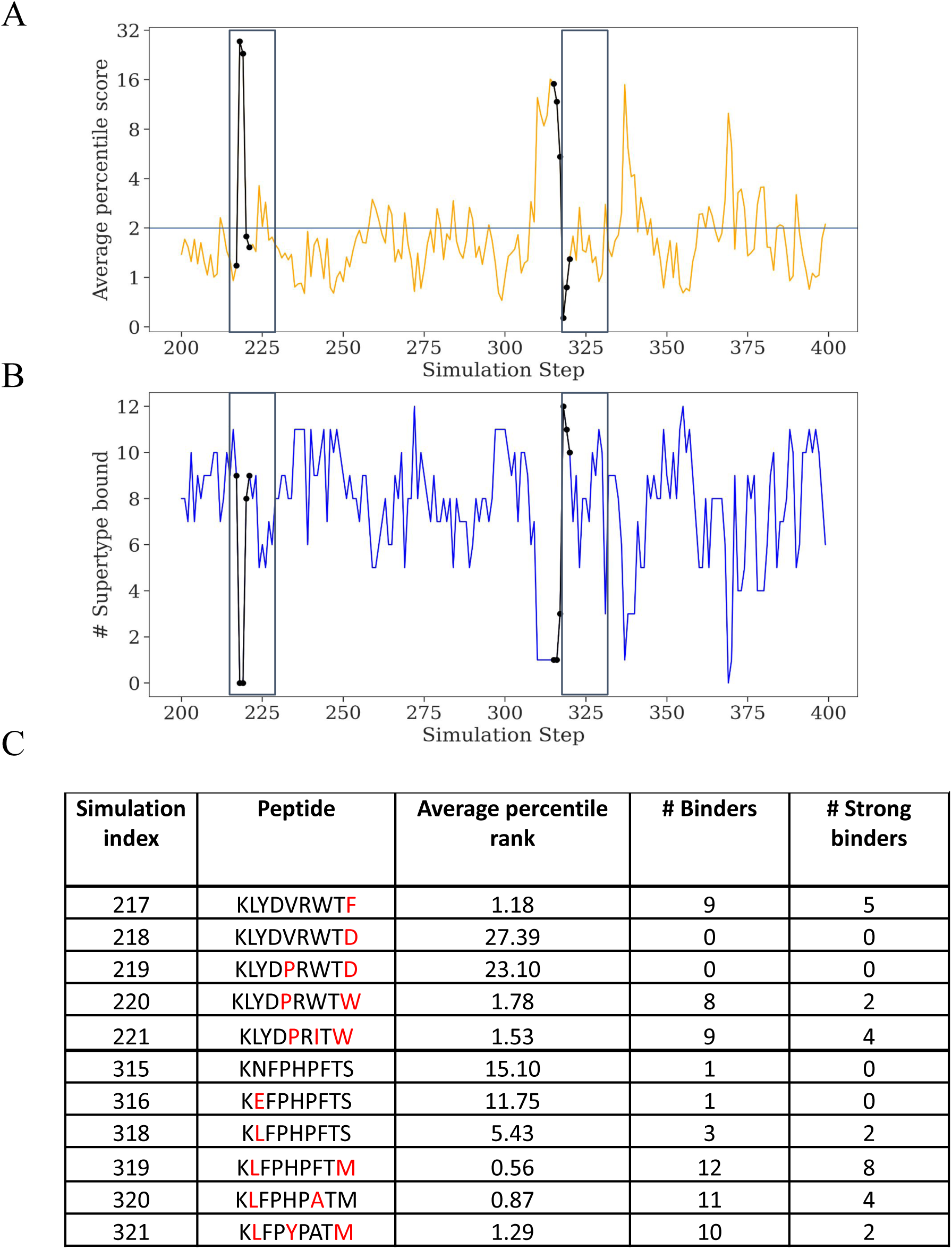
Visualization of super-HLA algorithm across 200 steps. (**A**) **Average percentile score across time.** The average HLA binding percentile rank across 12 representative alleles from all HLA class I supertypes is plotted as a function of simulation steps. Lower percentile rank corresponds to higher binding affinity. (**B**) **The number of bound supertypes across time.** For each simulation step, we compute the predicted number of bound supertypes using percentile cutoff of 2%, which is typically used to define HLA binders. In each plot, individual steps that sharply reduce the number of superbinders to avoid local maxima.were highlighted (217,315). A higher average percentile score corresponds to a lower number of bound supertypes. (**C**) A table of simulation steps 217-221 and 315-321 which correspond to the first black rectanbles marked in plots A and B. Each row denotes a single simulation step.

**Table 1:** List of supertype representative alleles used in the study.

| **Allele** | **Supertype** | **Allele Frequency\*** | **# of Binders in IEDB ^** |
| --- | --- | --- | --- |
| A*01:01 | A01 | 0.04843 | 22369 |
| A*02:01 | A02 | 0.15284 | 77727 |
| A*03:01 | A03 | 0.04272 | 12642 |
| A*24:02 | A24 | 0.18816 | 20660 |
| A*30:01 | A01A03 | 0.02505 | 4199 |
| B*07:02 | B07 | 0.04105 | 30084 |
| B*08:01 | B08 | 0.0296 | 9082 |
| B*27:05 | B27 | 0.01236 | 58349 |
| A*29:02 | A01A24 | 0.01626 | 8966 |
| B*40:01 | B44 | 0.0512 | 17936 |
| B*5801 | B58 | 0.02891 | 8127 |
| B*15:01 | B62 | 0.03431 | 15662 |

\* Solberg et al., Human immunology, 2008
^ Data extracted from IEDB on 17/4/2025

More formally, at each step, the algorithm randomly selects a position along the peptide *p*_*i*_ and introduces a random amino acid mutation at this position (**Algorithm 2.2.1-2.2.2**). The resulting mutated peptide is denoted by *p*_*i*_, which differs from *p*_*i*–1_by a single point mutation. We then compute the overall change in binding affinity between *b*_*hla*_*k*__(*p*_*i*_) and *b*_*hla*_*k*__( *p*_*i*–1_) defined by:

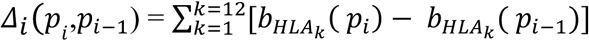

Where *b*_*HLA*_*k*__( *p*_*i*_) is netMHCpan4.1 percentile rank score

To accept or reject a given step, we used the following cost function, which was selected to allow the simulation to also accept steps that decrease the number of bound alleles with low probability, allowing it to explore additional regions in peptide space:

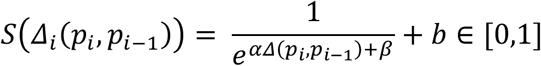

Specifically, we used *α* = 0.2, *β* = 2 and *b* = 0.02.

### Generating a superbinder pool using the *superHLA* algorithm

We ran ten simulations of the algorithm each starting the search procedure from a randomly generated peptide. Each simulation explored 40,000 unique peptides covering a total of 400,000 peptides. On average, each simulation generated ∼19,000 *superbinders* defined as peptides that bind 8-12 different supertype representatives. We computed the pairwise hamming distances between all pairs of putative superbinders obtained across the ten simulations and found that the median hamming distance between peptides was 7 and that the peptides did not cluster by simulation (**Figure 4A-B**). We found that the distributions of the number of superbinders bound by each peptide were similar across different simulations (**Figure 4C**). We also extracted all known experimentally verified superbinders from the Immune Epitope Database (IEDB)^19^. We found a total of 20 epitopes that bound 8 or 9 alleles from distinct supertypes, and one additional epitope that bound 10 alleles (**Figure 4D**). Utilizing the superHLA algorithm across the ten different simulations, we generated a set of 192,382 putative HLA superbinders i.e., peptides that bind to more than 8 different HLA representatives.

**Figure 4.**
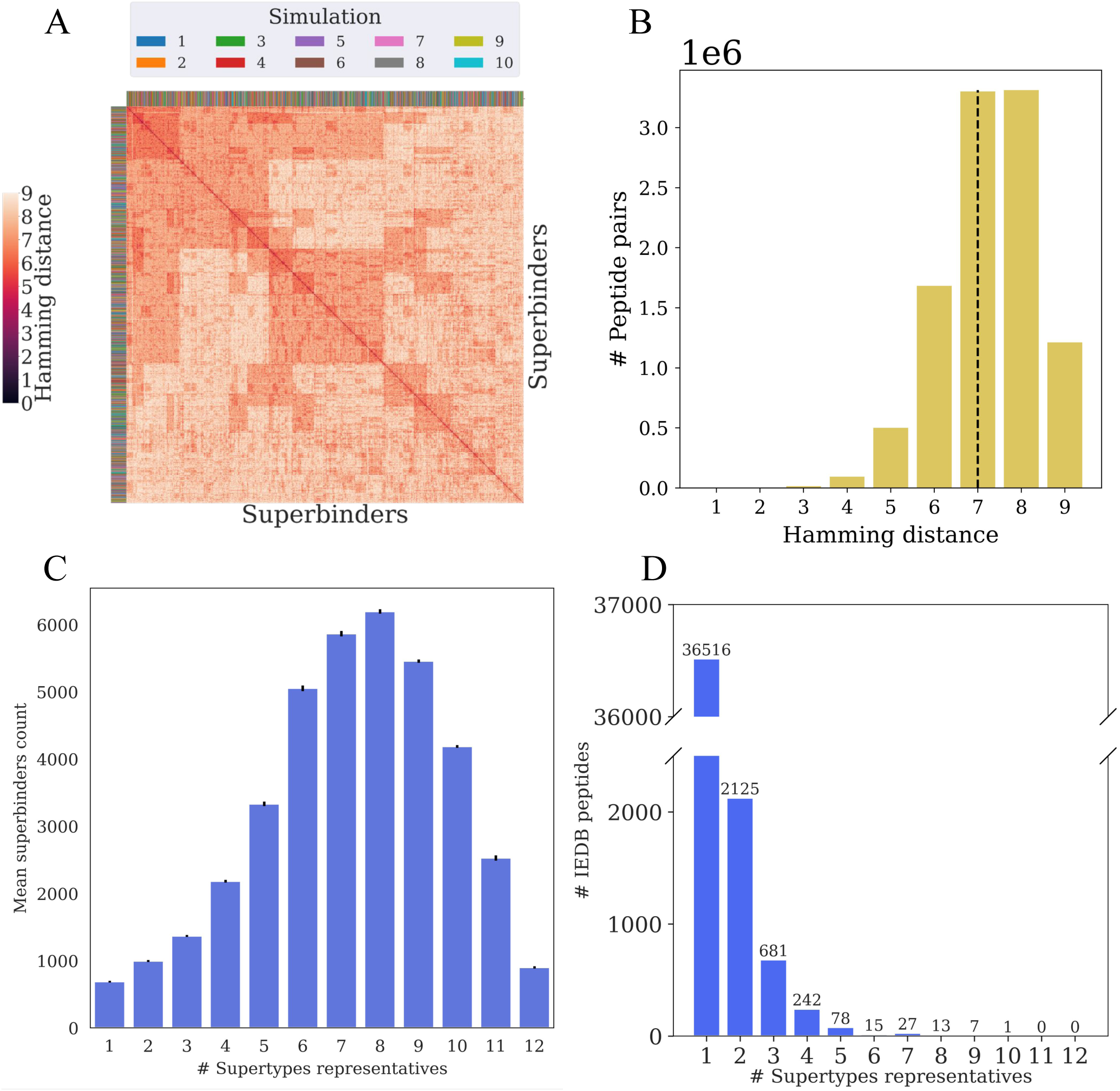
Comparison of the superbinder sets obtained from 10 independent runs of *super-HLA*. (**A**) Hamming distance matrix of the 4500 peptides generated using *superHLA*. Darker colors represent larger hamming distances. Peptides were clustered using linkage clustering. The color ribbons on both axes denote the run number from which each peptide originated. (**B**) A histogram of the pairwise Hamming distances between superbinders generated using *super-HLA* (n=4500). The median Hamming distance was 7. (**C**) A histogram of the average number of superbinder peptides obtained from 10 independent simulations each consisting of 40K simulation steps. Peptides were sorted by the number of supertypes they are predicted to bind (**D**) A histogram of the number of supertypes bound by experimentally verified 9-mer peptides in IEDB. We extracted all IEDB 9-mer peptides that were experimentally reported to bind to one or more HLA class I allele across the 12 representative alleles selected to represent each HLA supertype. Only 383 (1.05%) of the IEDB 9-mer peptides for these representative 12 alleles were reported to bind alleles from four or more HLA supertypes.

### superHLA generates superbinder peptides from a broad range of HLA supertype combinations

To further filter the list of putative superbinder peptides, we developed several filtering steps (**Figure 5**). We first grouped all 192,382 peptides into HLA binding preference groups, i.e., all peptides that bound a specific set of HLA alleles from the overall set of 12 alleles were grouped together. We found that out of the 784 possible HLA combinations of 8 or more alleles (each representing a unique supertype) we obtained putative peptides for 471 (60%) of these combinations. As a first filtering step, we clustered the peptides within each HLA combination using the CD-HIT clustering algorithm ^27^ ^28^. We chose a representative peptide from each CD-HIT cluster based on the binding potency of each peptide (see methods). In our initial simulations, we observed a strong bias towards amino acids P, D and E in position 4 of the peptides generated by our MCMC process. To reduce this potential bias, we opted to select representative peptides that include other amino acids at position 4 when possible. This first clustering step reduced our peptide set to 55,066 putative superbinders. To explore how the diversity of solutions within each of HLA combinations is affected by combination size (i.e. the number of supertypes in the group) we computed the cluster diversity ratio, i.e., the ratio between the number of unique clusters obtained by CD-HIT and the total number of peptides within this HLA combination (**Figure 6A**). As expected, the cluster diversity ratio was higher for smaller HLA allele combinations, likely because such combinations are less restricted by HLA binding preferences and thus allow for greater peptide diversity. However, there is also inherent variability in the number of unique solutions within a given HLA combination size (**Figure 6B**).

**Figure 5.**
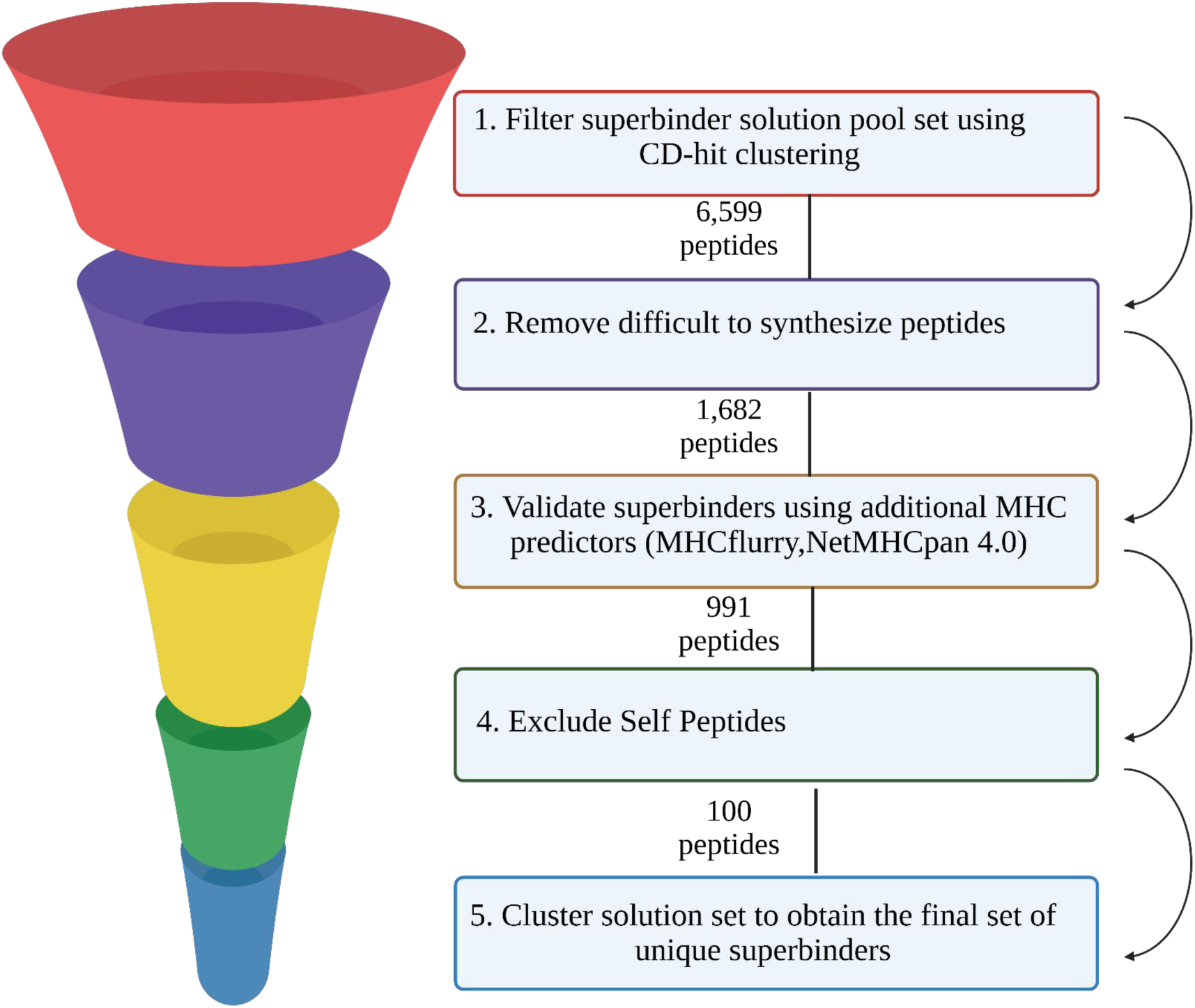
Filtering putative superbinders. Following the generation of a putative set of HLA superbinders, the following filtering steps were applied: (1) **Filter superbinder solution pool set using CD-hit clustering**. Two rounds of CD-hit clustering are used to filter the initial set of superbinders in order to obtain a diverse set of superbinders, yielding an initial set of 8435 superbinders. (**2) Removing difficult to synthesize peptides** - peptides that included a set of amino acid motifs that are difficult to synthesize such as ones containing GG were removed from the pool of peptides yielding a total of 6599 peptides. **(3) Validate superbinders using additional MHC predictors**. We computed the supertype specificity of each candidate peptide utilizing two additional HLA binding predictors - netMHCpan 4.0 and MHCflurry, and selected peptides that had broad binding profiles across all three predictors, yielding a set of 1682 peptides. **(4) Exclude self-peptides.** To eliminate self-peptides we compared the superbinder pool to the human peptidome and removed all peptides that were highly similar to any self peptide providing a total of 991 peptides. **(5) Cluster solution set to obtain the final set of unique superbinders.** A phylogenetic tree of all peptides was generated using linkage clustering was used to select 100 clusters from which a single representative peptide was selected.

**Figure 6.**
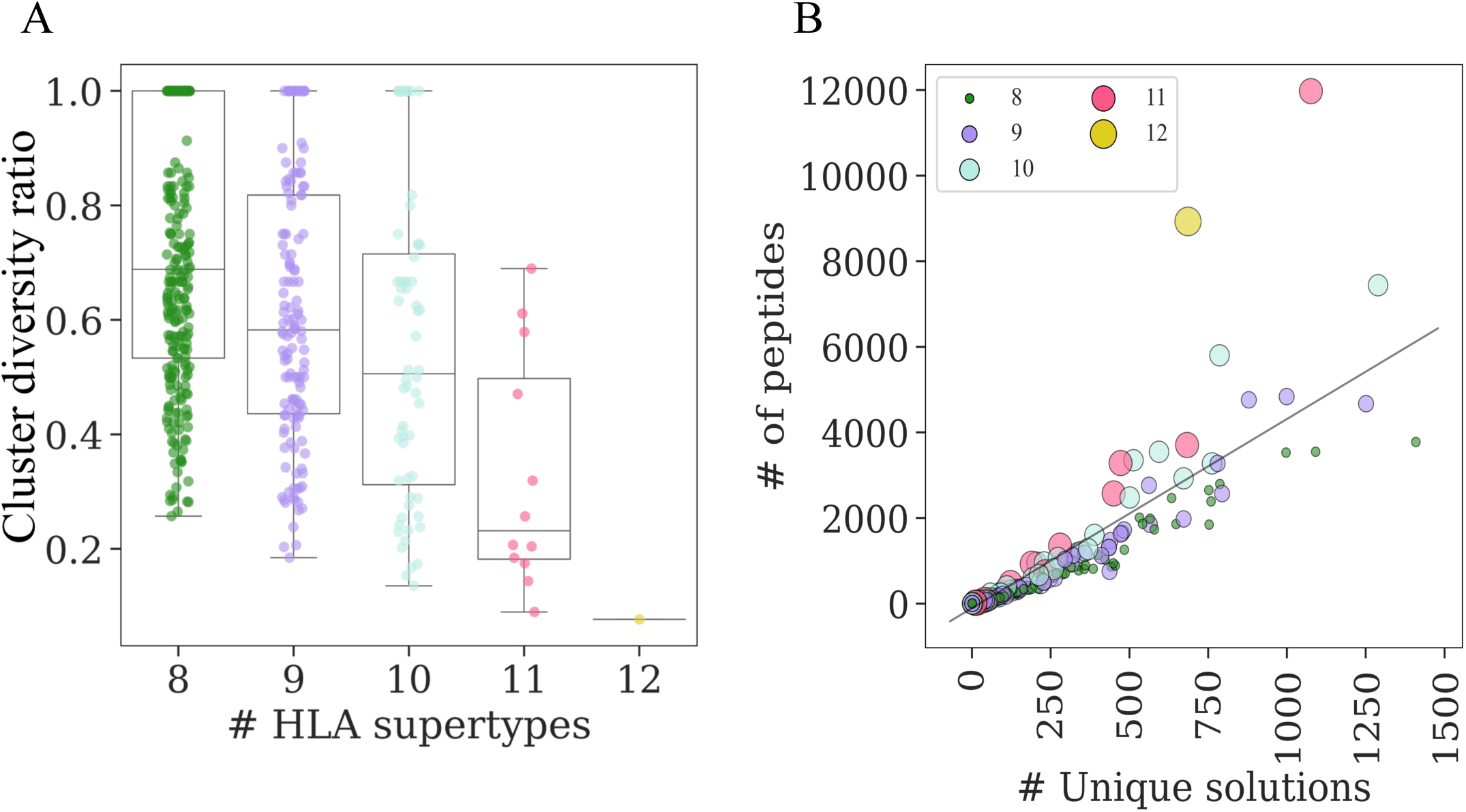
Superbinder diversity as a function of binding HLA allele combination sizes. We considered all sets of HLA combinations of size 8-12. For each combination, CD-HIT clustering was used to cluster all peptides that bound the specific combination of HLA alleles. **(A)** Cluster diversity ratios of specific clusters sorted by the number of unique supertypes their solutions bind **(B)** A scatter plot of the number of CD-HIT clusters (unique solutions) vs. cluster size for HLA combinations of 8-12 alleles spanning unique supertypes. Each unique HLA combination is colored by the number of alleles within that combination.

We then performed a second clustering step on the representative peptides from the first phase, combining solutions from all HLA combinations into a single set. Representatives were once more chosen based on their binding potency and by preferring amino acids other than P, D and E at position 4. Following these two clustering steps, a total of 8,435 peptides remained in our putative superbinder set. Each cluster is characterized by unique properties, including its binding motif and HLA allele combinations. Furthermore, all clusters contained solutions from an average of 3 simulations (**Figure 7**). The remaining set of peptides were then subjected to several additional filtering steps which included removal of peptides that are difficult to synthesize, exclusion of self peptides and the use of additional binding predictors to select only peptides that are predicted to bind based on multiple predictors (**Figure 5** and methods). To cross-check our putative superbinder set, we utilized two additional MHC binding predictors (neMHCpan 4.0, MHCflurry) to predict the binding affinity of each of the peptides in our set for all 12 alleles^29,30,31^ (see methods). We selected the 3000 top ranked peptides from each of the three predictors and then chose their intersection (**Figure 8**), resulting in a set of 1,682 putative superbinders. To obtain the final set of unique putative superbinders, we used a maximum likelihood phylogenetic-based clustering algorithm^32^ to obtain 100 clusters and selected a representative peptide from each cluster using the max cluster criterion (see methods).

**Figure 7.**
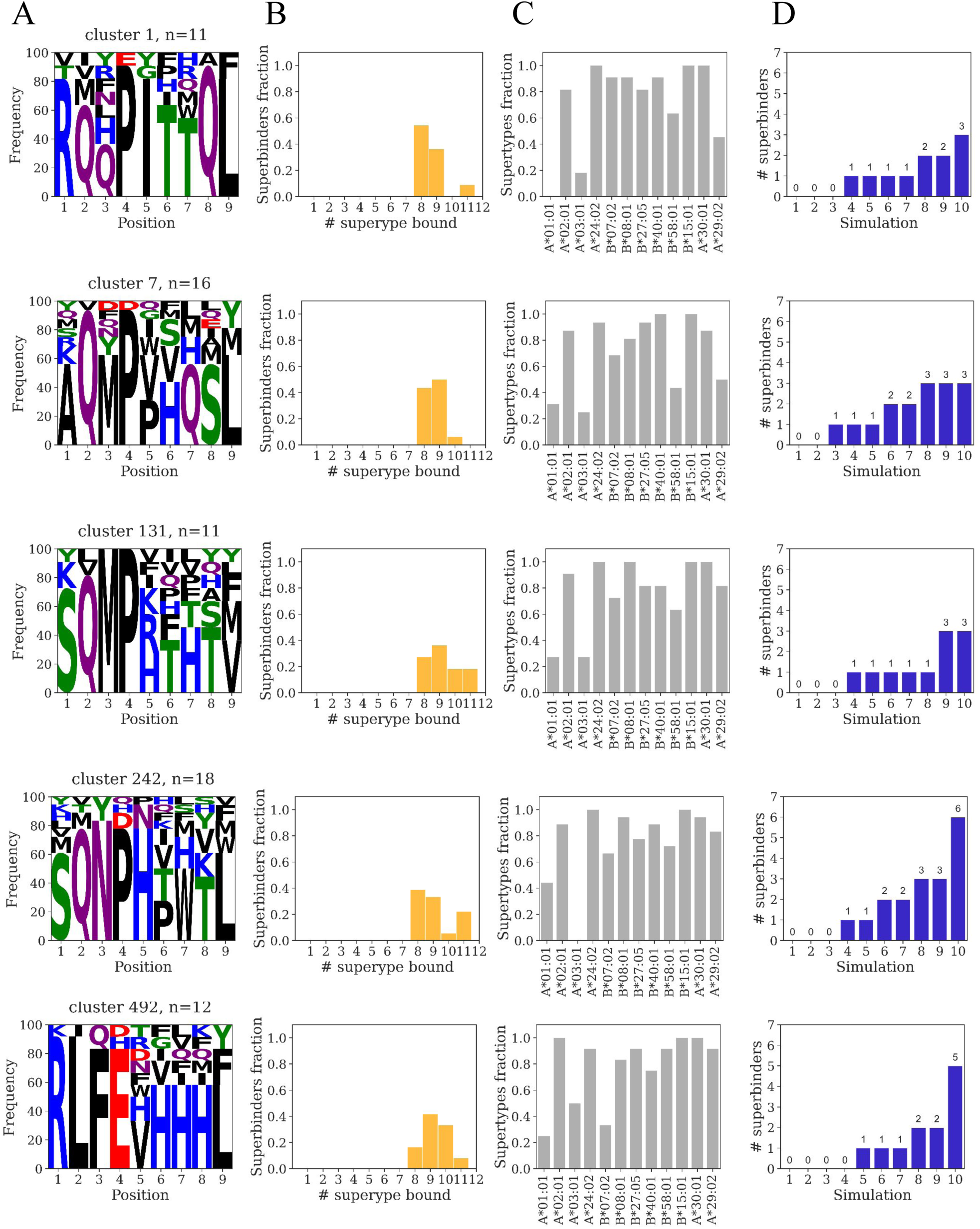
Properties of representative clusters of superbinders. Five representative clusters are presented, obtained following two rounds of CD-HIT clustering (see methods). (**A**) Logo plots of the distribution of amino acids in each position of the peptides within the cluter. (**B**) A histogram of the number of peptides within a cluster sorted by number of supertypes they bind; (**C**) A histogram of the frequency of peptides within the cluster that bind each of the representative allele spanning the 12 HLA class I supertypes (**D**) A histogram of the number of peptides within the cluster obtained in each of the 10 *super-HLA* runs used to generate the initial superbinder peptide pool.

**Figure 8.**
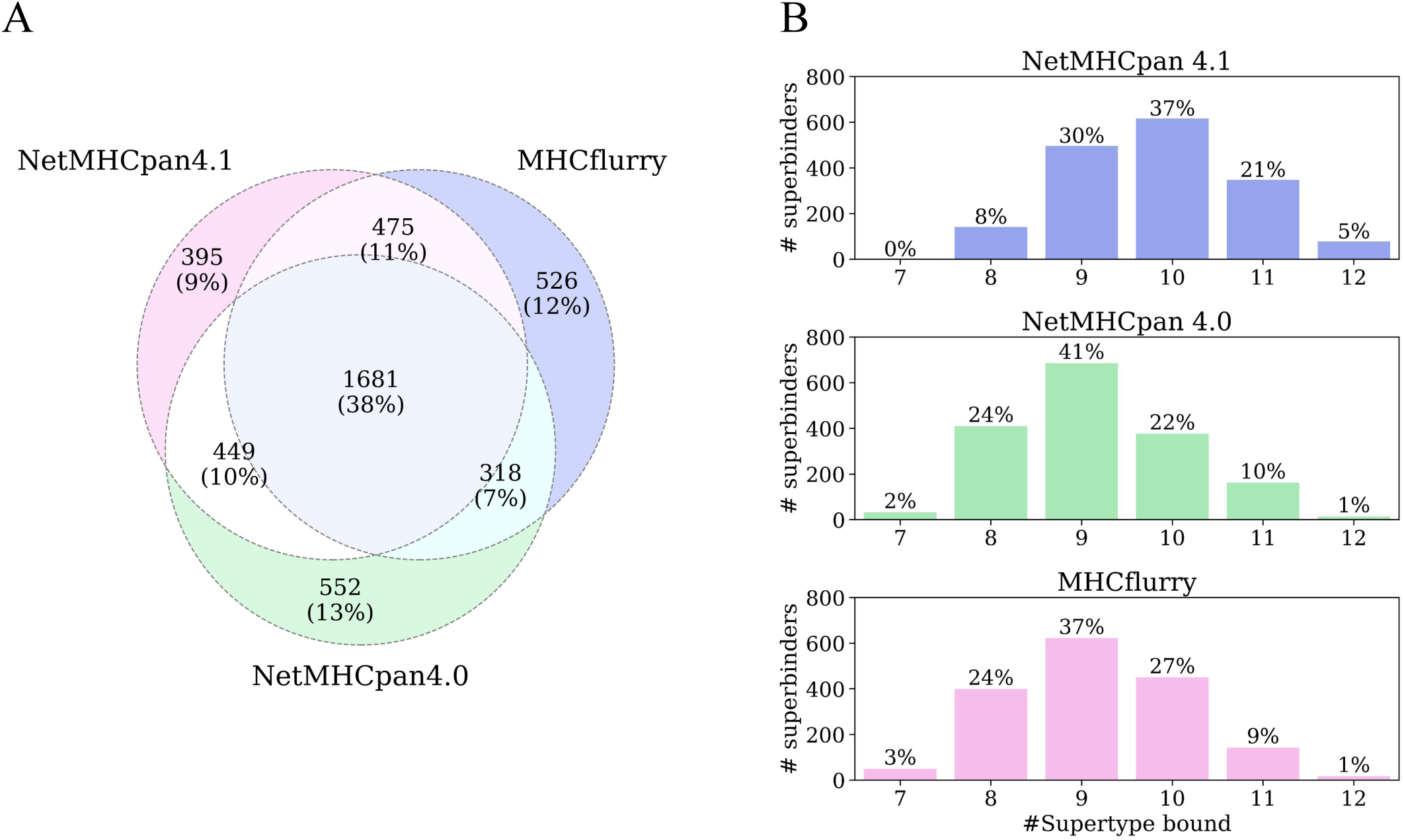
Filtering putative superbinders using two additional MHC prediction methods. We used netMHCpan 4.0 and MHCflurry to further filter our list of peptides considering peptides that were highly ranked across all three predictors. Each predictor was used to select a subset of peptides from the putative set of superbinders generated by the superHLA algorithm. Using each predictor, we computed the average percentile rank for the 8 alleles with the lowest percentile ranks. Using this score we selected 3,000 top ranked peptides from each of the three predictors, and then chose the intersecting peptides. **(A)** Venn diagram of the number of peptides that were identified by each of the three binding predictors used. A set of 1681 (38%) peptides were highly ranked by all three prediction methods**. (B)** Histograms of the number of supertypes bound by the set of 1681 peptides that were selected based on all three binding prediction methods.

To explore the diversity of the 100 superbinders selected we created a position-specific scoring matrix (PSSM) of the amino acids across all 9 positions of the peptides (**Figure 9A**). The hamming distances between all pairs of 100 peptides ranged from 4 to 9 with a median of 7 (**Figure 9B**). We observed that the amino acid distribution at positions 2 and 9 - known MHC class I anchor residues - was biased toward non-polar amino acids. Furthermore, position 4 showed a strong bias toward Proline, Aspartate, and Glutamate (P, D, and E). (**Figure 9C-D**). We therefore opted to select a representative peptide from each cluster by selecting the peptide with the maximal binding affinity to all 12 supertypes and when possible, selecting a representative that did not include P, D or E at position 4 of the peptide.

**Figure 9.**
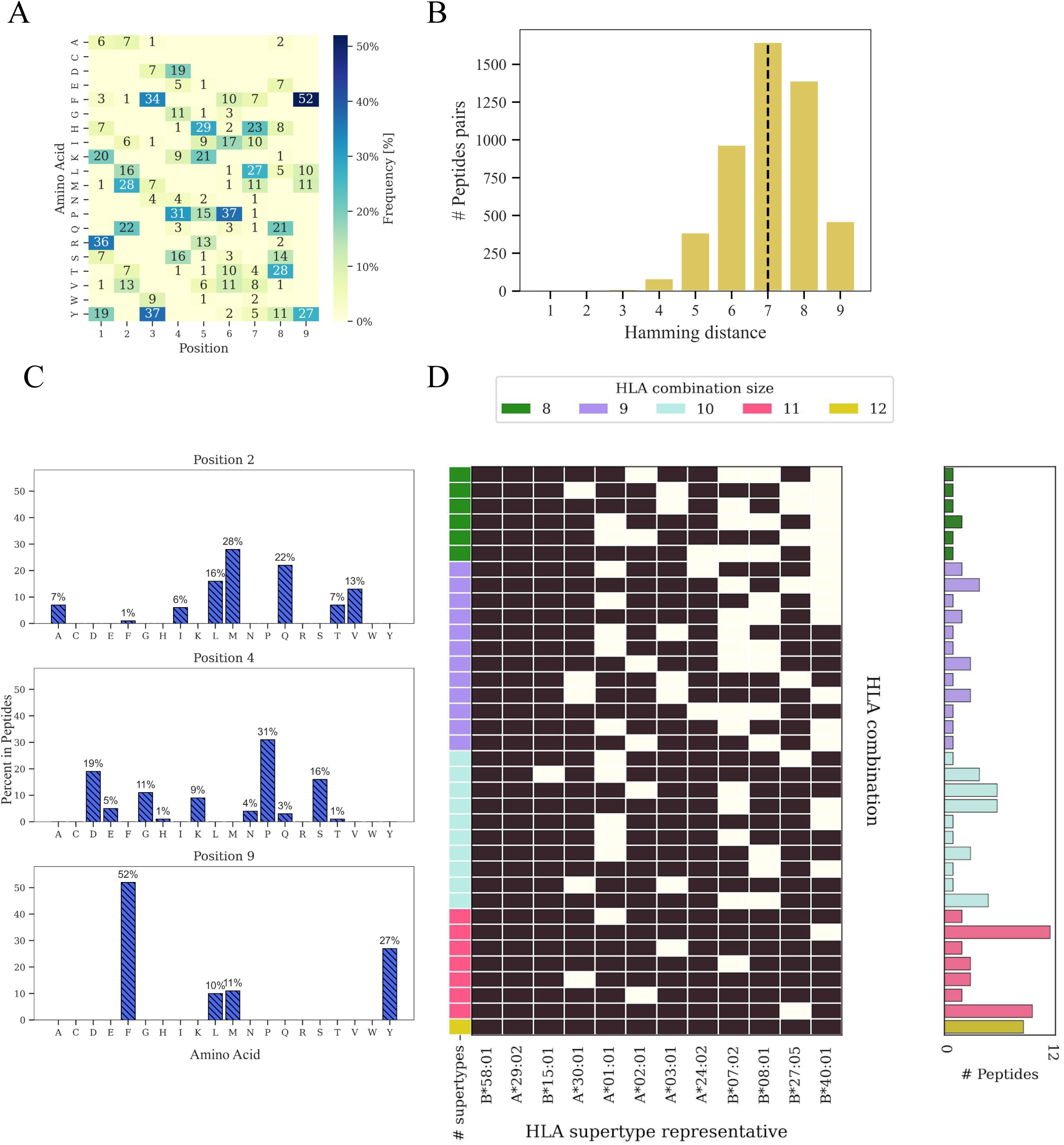
Properties of the 100 putative superbinder peptides selected for experimental validation. **(A)** Amino acid frequencies across all 9 positions of the final filtered superbinder set. Numbers represent the number of peptides containing a specific amino acid at a given position. **(B)** A typical histogram of pairwise hamming distances from one of the simulations performed (**C**) distributions of amino acids at positions 2, 4 and 9. Positions 2 and 9 are classical anchor residues for HLA class I binding. Position 4 has an apparent preference for proline, aspartate and glutamate in the initial pool of predicted superbinders. **(D)** The set of HLA combinations of size 8–12 covered by the superbinder validation set is shown. HLA combinations are ordered by the number of alleles bound, ranging from 8 to 12. The histogram on the right displays the number of unique peptides identified for each HLA class I allele combination. The color of each bar indicates the number of HLA alleles predicted to bind to each superbinder.

We then grouped peptides by the specific combinations of HLAs that they cover and found that some combinations are covered by multiple peptides, while others are only covered by a single peptide (**Figure 9D**). To further explore the sequence similarity within the final superbinder set, we identified all matching k-mers of size 4-6 between all pairs of peptides (**Supplementary Figure 1**). No shared 6-mer sequences were found, and only a single 5-mer peptide (RMFGP) was observed in two specific peptides (“RMFGPPLTF”, “RMFGPVQTY”). In contrast, 33 distinct 4-mer sequences were found to occur in two or more peptides.

### Experimental validation using HLA binding assays

To experimentally validate our final list of putative superbinders, we synthesized the 100 selected peptides generated by *superHLA* and tested their binding to a representative set of HLA alleles from the 12 supertypes **(Table 3)** using biochemical HLA binding assays^33,34^. In these assays, the binding affinity of a given peptide to a specific HLA allele is measured using a competition assay with a reference peptide^35^. Validation was performed in two steps: First, all 100 putative superbinders were tested against 6 HLA alleles from 6 supertypes: A*01:01, A*02:01, A*03:01, A*24:02, B*07:02 and B*40:01. Following this step, we selected all peptides (n=24) that were found to bind two or more supertypes from this set. These 24 peptides were then further tested against the remaining 6 supertypes: A*29:02, A*30:01, B*08:01, B*15:01, B*27:05, B*58:01.

**Table 2:**
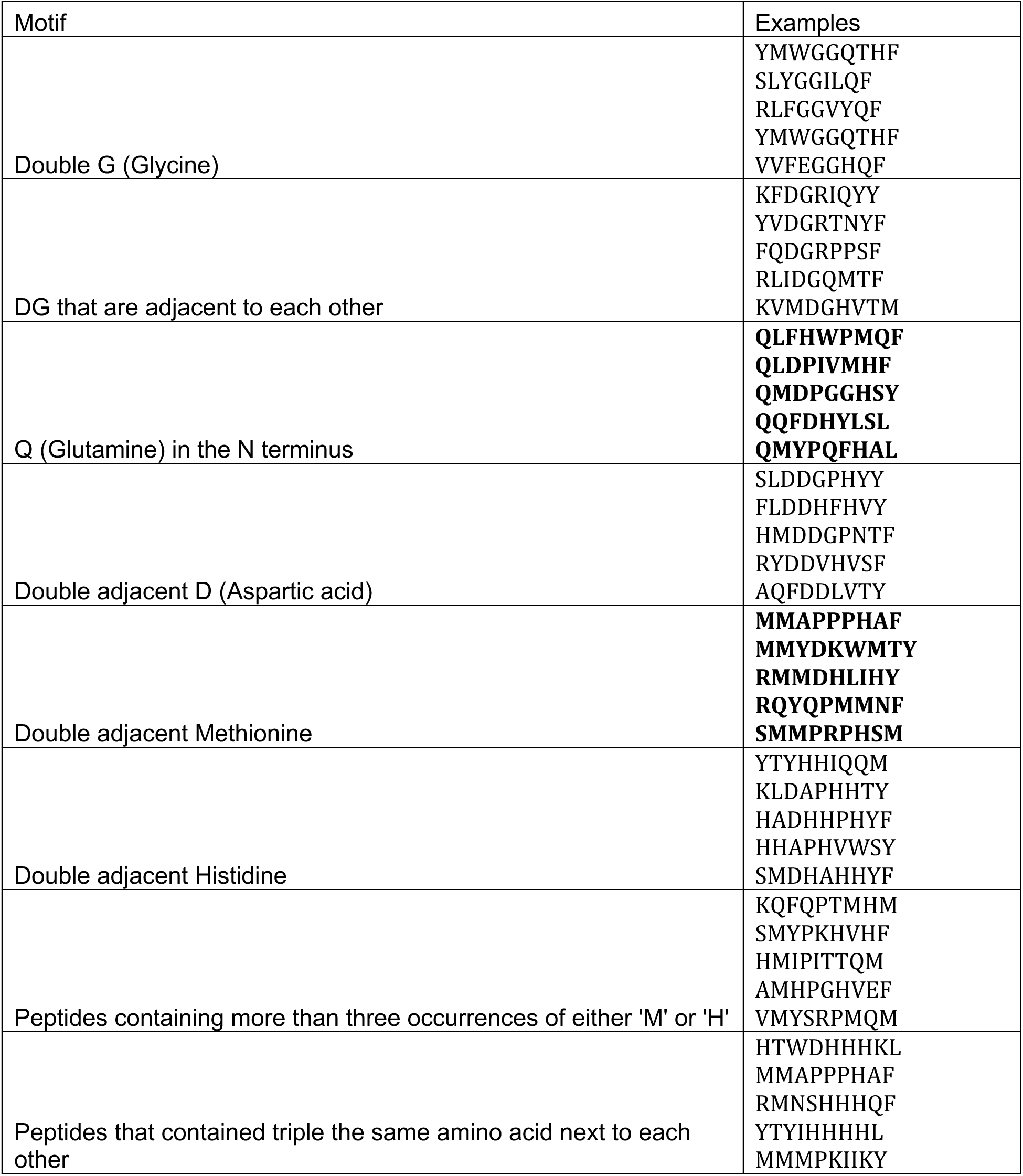
Examples of peptides with motifs that were removed from our solution set due to difficulty of synthesis.

**Table 3:**
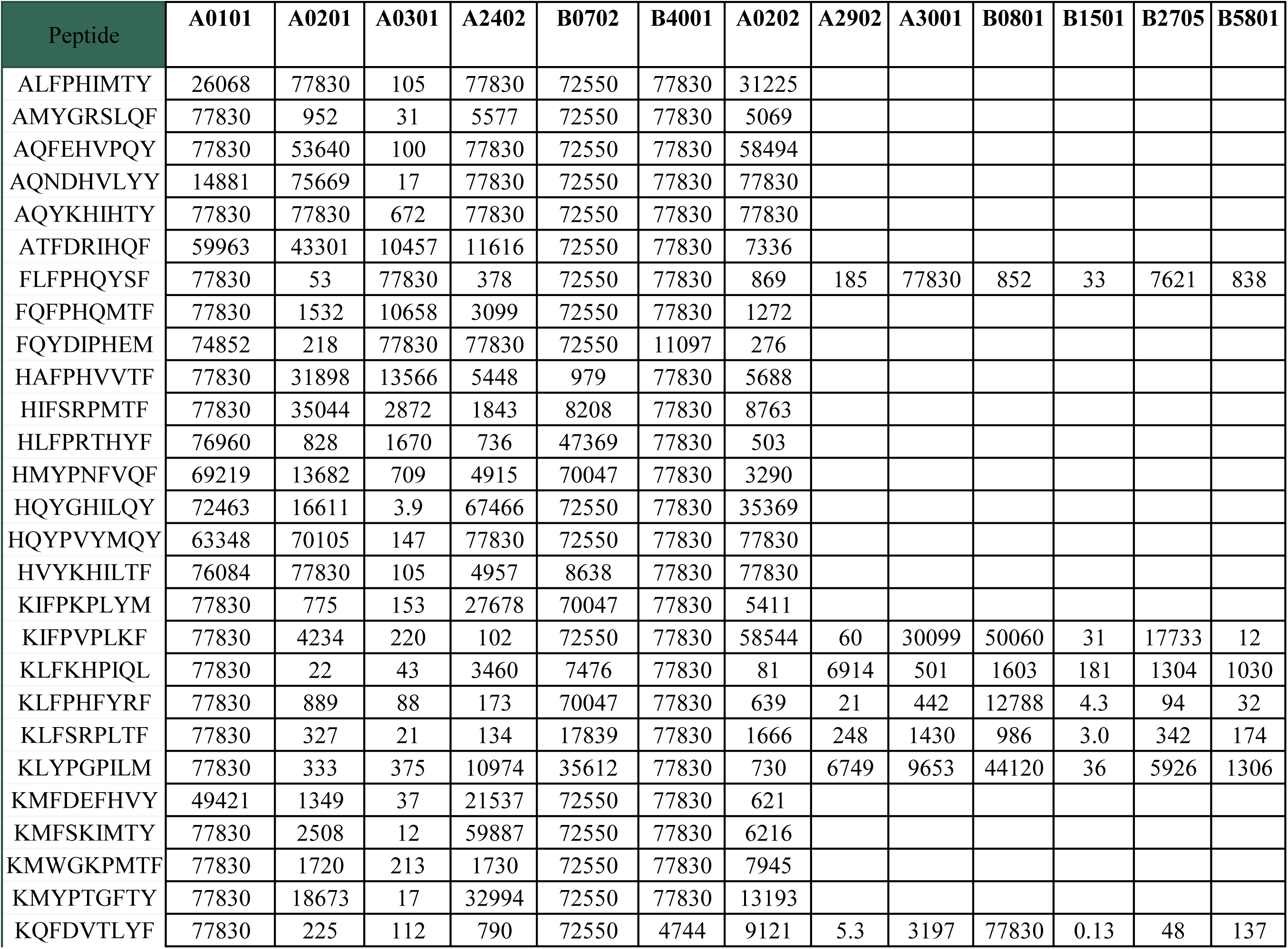
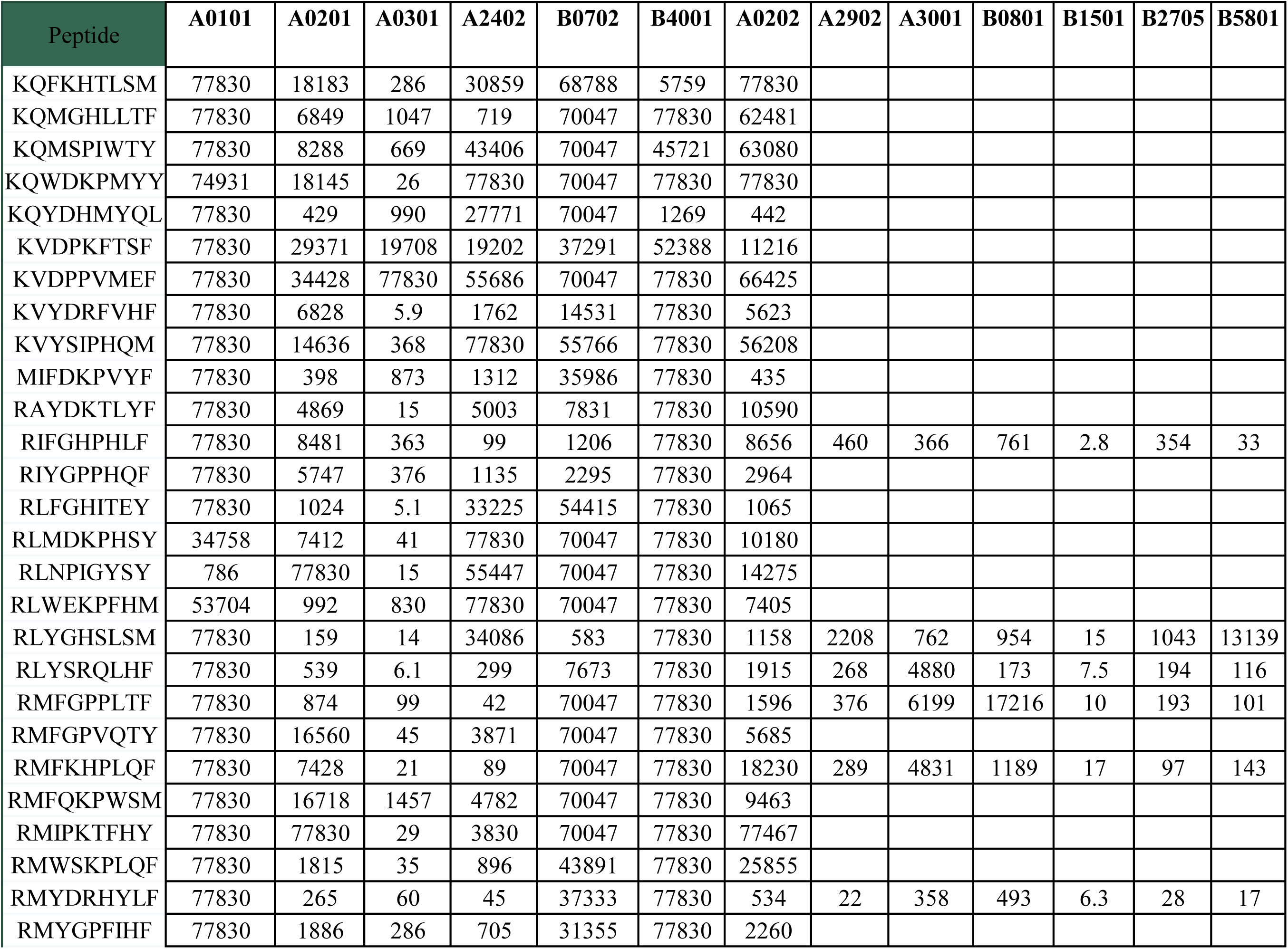
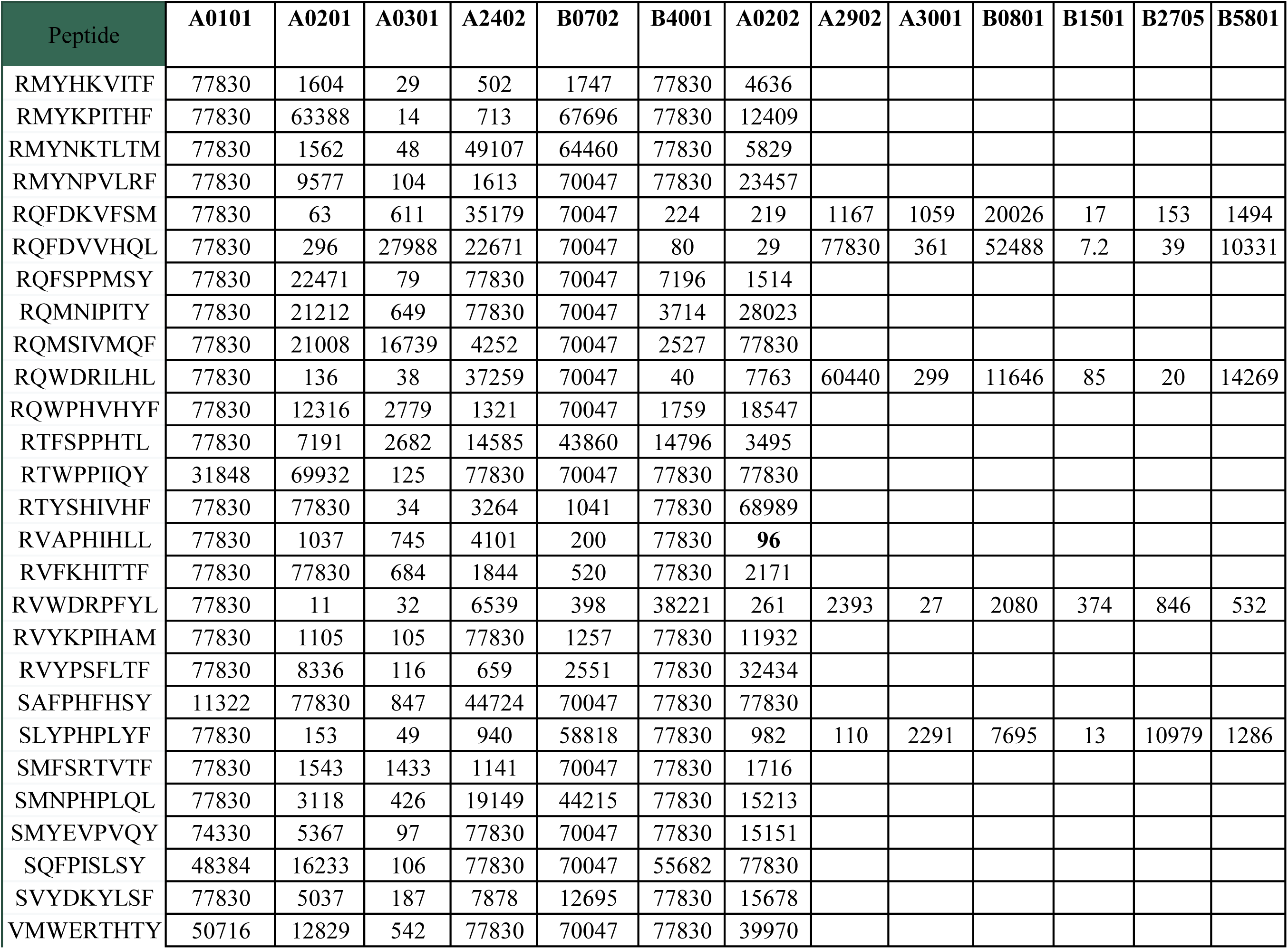
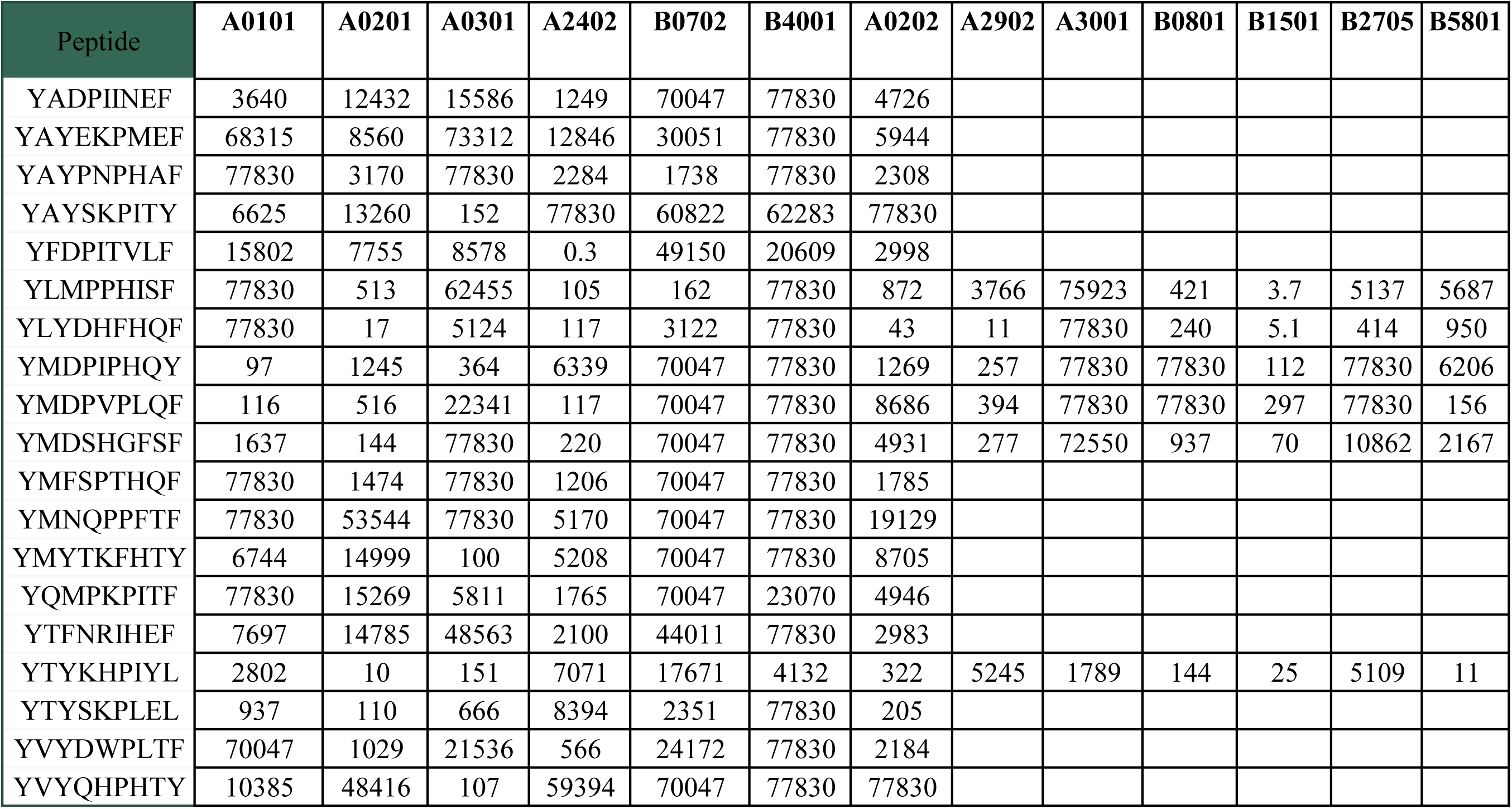
IC50 binding of the experimentally validated peptides.

We found that 21 (87.5%) of the 24 peptides tested against all 12 supertypes, bound 4 or more supertypes (**Figure 10A**). One peptide (RMYDRHYLF) bound alleles from 9 supertypes and four more bound alleles from 7 supertypes. All 24 peptides bound the B*15:01 allele and only two peptides bound the A*01:01 allele (**Figure 10B**). Since not all putative superbinders were predicted to bind to all 12 supertypes, we also computed the recall (true positive rate) of predictions for each allele. We found that the recall ranged between 2.86% (A*01:01) to 100% (B*15:01) with an average recall of 36.24% (**Figure 10C**).

**Figure 10.**
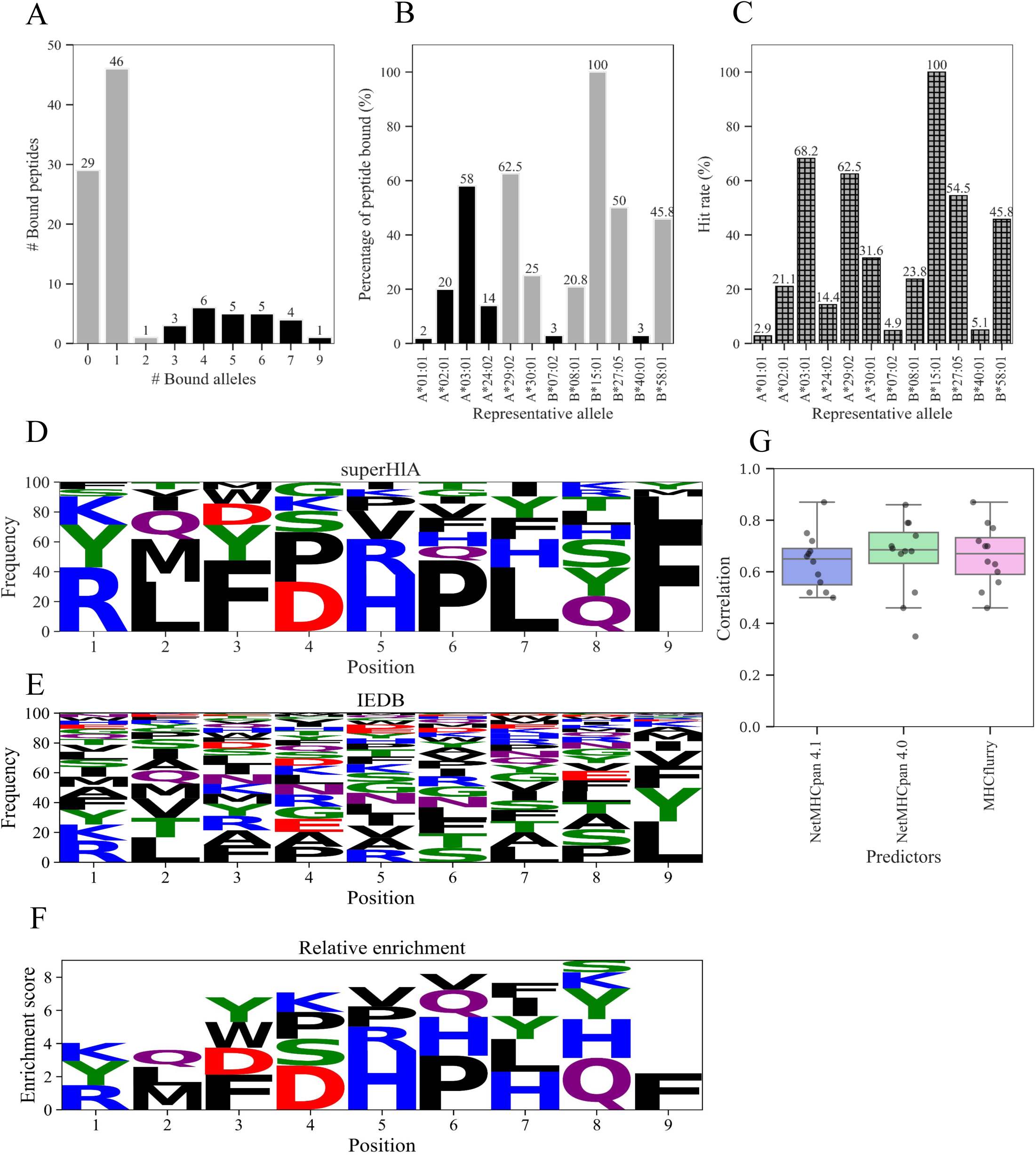
Properties of the experimentally verified superbinders. **(A)** HLA binding assays were conducted in two phases. In the first phase, all 100 peptides were tested across 6 HLA alleles representing six unique supertypes. Peptides that showed binding to three or more alleles in this set (n=24, orange bars) were then tested against an additional set of 6 HLA alleles representing the remaining six supertypes. Of the 24 peptides tested in the second round, 21 (87%) were found to bind at least one additional HLA allele from a different supertype. **(B)** Percentage of superbinder peptides bound to the representative HLA allele from each supertype. Grey bars represents HLAs that were tested on all putative superbinders while black bars bars denote the HLA representatives that were tested across part of the putative superbinders (n=24) **(C)** Histogram of recall rates in the experimental validation set per allele. **(D)** Logo plot of peptides that bound 4 or more different representative alleles in our validation set (n=21). **(E)** Logo plot of peptides that bound 4 or more representative alleles in IEDB (n=383). **(F)** Relative Enrichment for experimentally verified peptides in our set vs. those in IEDB. **(G)** Boxplots of the correlation between the predicted binding percentile ranks and the experimentally measured binding scores for experimentally validated superbinders. Each marker represents a single HLA allele..

We then generated sequence logo plots for the 21 experimentally verified superbinders and compared them to the sequence logo plot of all experimentally reported 9 aa long superbinders in the IEDB database (**Figure 10 D-F**). We found that the *superHLA* superbinders were enriched for hydrophobic residues at position 2 (L or M), basic residues at position 5 (H or R) and hydrobphobic residues at position 9 (F, L or M). These preferences were markedly different from those of the IEDB superbinder peptides (**Figure 10F**).

Comparing the experimentally measured IC50 binding values of the 24 putative superbinders to their predicted percentile score of the HLA binding predictors, we found correlations ranging between 0.5 (B*27:05) to 0.87 (A*24:02) for netMHCpan 4.1 (**Supplementary Figure 2A**). Interestingly, the correlations with netMHCpan 4.0 percentile scores were higher for all alleles except for four alleles (B*27:05, A*03:01, A*24:02 and B*40:01, **Supplementary Figure 2B-C**), as well as those of MHCflurry (**Supplementary Figure 2C**). Overall, peptides predicted to bind to a larger set of HLA supertypes, also bound more experimentally verified alleles, but these numbers were typically 3-6 fewer supertypes than predicted (**Supplementary Figure 3**).

## Discussion

Here we studied HLA superbinders – 9-mer peptides that bind to multiple HLA alleles from different HLA supertypes. To systematically study these peptides we developed *super-HLA* - a novel *in-silico* method for generating HLA class I superbinders, peptides that could be presented by a diverse set of HLA alleles ^18^. Our method uses an MCMC optimization process in order to sample the peptide space, taking into account 12 representative HLA class I alleles from all 12 supertypes. We defined superbinders as peptides that bound 4 or more representative alleles.

We found that despite the extensive diversity of HLA alleles, experimental data, as well as our simulations demonstrate the existence of superbinders. In our analysis of the Immune Epitope Database (IEDB) we found that only 1% of all experimentally verified binders are superbinders, most of which were identified through peptide elution measurements. Only a single peptide was found to bind 10 HLA alleles, and just seven peptides bound 9 alleles. Using *super-HLA*, we were able to identify 21 superbinders out of 100 tested peptides. Since only 24/100 peptides were tested against a representative HLA from all 12 supertypes, it is likely that there are additional superbinders within this set.

We found that superbinder solutions tended to converge to specific regions of the peptide space, as evidenced by peptide clusters that often included peptides from multiple independent simulation runs. To select the final set of 100 peptides for experimental validation, we applied several filtering steps, including the selection of representative superbinders that were diverse in sequence space. Starting from a putative set of 192,382 peptides, this process resulted in a final list of 100 candidates selected for validation. It is therefore highly likely that additional superbinders - similar to those experimentally verified by *super-HLA – that* were filtered out using our selection strategy. Taken together, our analysis suggests that superbinders are more prevalent than previously reported.

We found that while our method was able to generate a large and diverse set of superbinders, there were specific recurring binding motifs within superbinders. We also found some HLA allele combinations for which superbinders were much more readily obtained. Specifically, at some positions, our superbinders show a significant bias toward particular amino acids. Charged amino acids were observed in the N-terminus (R, K, H, S, Y). Positions 2 and 9, corresponding to the B and F binding pockets of the HLA binding groove and largely dictate its peptide specificities, exhibited high selectivity. In both of these positions there was a tendency toward non-polar amino acids (L, M and F) which was more dominant in position 9. Position 2 showed greater variability in the characteristics of the possible amino acids, and also included polar amino acids (Q, S, T). We also found that in some cases a single mutation in the anchor residues 2 and 9 could result in increased binding affinity to 8 representative HLA alleles. Interestingly, our analysis also identified specificity at position 4 of the peptide, which mostly included P, D and E, with P being the most frequently observed amino acid within the superbinder set. Proline is the only amino acid where the side chain is connected to the protein backbone twice, leading to conformational rigidity. We therefore computed the frequency of proline in position 4 of all known epitopes in IEDB and PDB. We found that 7.2% of experimentally verified 9-mer peptides that bind HLA class I alleles contained a proline in position 4. However, only four of these were superbinders (QQRPDLILV, FVMPIFEQI, MTFPLHFRS and FFSPFFFSL). As it is currently unclear whether this bias reflects a true property of HLA class I superbinders, or is simply some artifact of the HLA binding predictors, we significantly reduced the number of peptides that contain P, D or E at position 4 in our experimental validation set.

The prediction of MHC-peptide binding using motif matrices and sequence motifs assumes that peptide residues contribute independently to HLA binding. Such an assumption, in general, is well supported by experimental data, but there is also evidence indicating that the contribution of peptide residues to MHC-binding is influenced by neighboring residues^36^. Indeed, when we analyzed the k-mer distributions of putative superbinders, we found specific recurring motifs, mostly of size k=4 including PLTF (positions 6-9), PHYF (positions 6-9), PHTF (positions 6-9), FSHP (positions 3-6) and YSHP (positions 3-6)). Each of these 4-mers appeared in different HLA combination sets, suggesting that specific k-mers may be associated with superbinders that bind to specific subsets of the 12 representative HLA alleles. Interestingly, we found 2 peptides in our set of 21 validates superbinders that included the PLTF motif.

We also found that there were specific combinations of HLA alleles that were more frequently obtained in our simulations, while other combinations were not observed at all. For example, only 471 out of the 794 possible HLA combinations of 8 or more alleles were observed in our putative superbinder set. As expected, the peptide diversity was higher for smaller HLA allele combinations, but there is also inherent variability in the distribution of the superbinder diversity score within a given HLA combination size. In the final list of the 100 final superbinders 8/495 (1.6%) of the 8 HLA allele combinations were observed, 12/220 (5.5%) of 9 HLA alleles combinations included, 12/66 (18.2%) of 10 HLA alleles combinations included 7/12 (58.3 %) of the 11 HLA allele combinations were observed, and all possible combinations of 12 alleles were sampled. After filtering the superbinders using the various filtering steps utilized by *super-HLA* we found that the most favored combination of HLA alleles included 11 HLA representative alleles, excluding HLA-A*01:01, which included peptides from different simulations, and was characterized by a preference of negatively charged amino acids in position 1 and 5.

Previous studies have found that peptides binding with higher affinity to MHC are more likely to be displayed on the cell surface where they can be recognized by T lymphocytes^37^. *Super-HLA* builds upon this to generate peptides that are highly likely to bind to multiple HLA alleles. However, there are other factors that govern the immunodominance of a specific peptide, which include cleavage propensity and the expression level of a given antigen within the antigen presenting cells. Since super-HLA generates de-novo putative superbinders, it does not factor these into its optimization process.

While peptide binding to MHC molecules is the key feature in cell-mediated immunity^38^, a peptide that binds a specific HLA allele will not necessarily become a T-cell epitope, as T-cell immunogenicity is contingent on several additional factors including the stability of the peptide-MHC complex, TCR specificity and abundance, the phenotype of the responding T cell, and presence or absence of secondary signals^39^. Therefore, future work is needed to assess if our superbinder peptides may indeed be T-cell epitopes.

There are several potential applications of HLA superbinders. First, they may serve as potent adjuvants to antibody based immunotherapies, similar to current approaches that rely on using dominant immunogenic epitopes such as the CMV pp65 epitope^40^,^41^. The advantage of using a superbinder is that it can stimulate T-cells in individuals carrying multiple dominant HLA alleles. Another potential application of superbinders may be as potent adjuvants for vaccines. Previous studies have reported the use of a 13aa sequence that was used to enhance CD4 responses^42^. Another study reported that several peptides on the NY-ES-1O cancer-testis antigen bound multiple HLA class II alleles and elicited CD4+ T-cell responses^43,44^. However, a key limitation of our current approach is that the superbinders generated by super-HLA are synthetic and were not previously observed by the immune system. Therefore, peptide vaccination will be required to elicit T-cell responses against these peptides. Given recent advances in our ability to generate potent immune responses against peptides using such vaccines^45^,^46^, this hurdle may be overcome.

An important advantage of the Super-HLA framework is its flexibility and adaptability to a wide range of use cases. While our study focused on identifying peptides that bind to a predefined set of 12 representative HLA class I alleles from distinct supertypes, the method can be readily extended to target any custom combination of HLA alleles. This makes Super-HLA suitable for personalized applications, such as tailoring peptides for specific population HLA frequencies or for individual HLA genotypes in clinical contexts. In addition, the MCMC-based optimization can be initialized from a peptide of interest rather than a random starting point. This allows the algorithm to explore the local peptide neighborhood and identify sequence variants that preserve or improve multi-HLA binding potential. Moreover, Super-HLA can incorporate additional constraints into the optimization process to restrict sampling to specific regions of the peptide space. For example, constraints on biochemical properties (e.g., charge, hydrophobicity), avoidance or preference of specific motifs. This enables focused exploration of peptide subspaces with enhanced relevance to specific biological, immunological, or clinical objectives. Together, these features make Super-HLA a versatile tool for the rational design of broadly presented peptides, supporting applications in vaccine development, immunotherapy, and basic immunological research.

Our current work also suggests that a similar approach could be utilized to develop CD4^+^ superbinders, which may be important for providing potent T-cell help for antibody-based vaccines. However, additional work is required, due to the differences in binding preferences of HLA class II alleles as compared to HLA class I alleles, which we have utilized here. There are several other possible further enhancements of our method: first, we may also take into account peptide cleavage motifs to improve the presentation potential of our HLA superbinders. Secondly, we may also consider longer peptides with nested binding motifs, which may provide overlapping motifs. These are more representative of our current knowledge of T-cell epitope hotspots, which have been extensively studied in the context of HIV infection and vaccination^16,17,47^.

In summary, here we presented a novel algorithm for de-novo generation of putative superbinders. Using this algorithm to identify 100 putative superbinders, we showed that 21 of them bound 4 or more alleles from distinct supertypes. Unlike naturally occurring immunodominance hotspots, which typically encompass overlapping peptide motifs that bind multiple HLA alelles, our work demonstrates that HLA superbinders are much more prevalent than previously appreciated, and identifies specific regions in peptide space that include such immunodominant peptides. Our work provides a first step and necessary step for designing highly immunogenic peptides that may be incorporated into future vaccines and T-cell based therapies.

## Methods

### HLA binding prediction

For a given peptide, we used netMHCpan 4.1 - an HLA binding prediction algorithm - to predict the binding affinity to the set of 12 alleles representing all 12 HLA class I supertypes **(Table 3)**. The algorithm outputs a percentile rank for a given peptide, where lower ranks correspond to higher binding affinity. Typically, a 2% cutoff is used to define a binding peptide (corresponding to a binding affinity < 500nM), and a 0.5% cutoff is used to define a strong binder (corresponding to a binding affinity < 50nM).

### Clustering peptides by sequence similarity

To filter our putative superbinder list we utilized the CD-HIT clustering program ^28^. Briefly, CD-HIT is an algorithm that uses a greedy incremental clustering method. It produces a set of closely related sequences (proteins or nucleotides) from a given fasta sequence file. When all sequences are of equal length - as is the case in our application - the algorithm greedily assigns the first sequence as the first cluster representative. Each subsequent sequence is either added to an existing cluster or used to start a new cluster if it doesn’t meet the similarity threshold with any existing cluster representative^27,48^. Two rounds of CD-HIT clustering were performed to address limitations arising from its greedy, incremental clustering strategy, which can occasionally place highly similar peptides into separate clusters. In the first round, we applied a high-resolution clustering step over each group of peptides sharing the same HLA binding preferences. For each cluster we selected a representative peptide based on the average percentile rank across 8 HLA alleles with the lowest percentile ranks. In the second round, we clustered the representative sequences from the initial round, allowing peptides with different binding profiles—but high sequence similarity— to be grouped together. Representative sequence for each cluster in the second round were selected as in the first round, but also excluded any peptides that had P, D or E in position 4 of the peptide. This iterative approach enabled us to consolidate redundant solutions and produce a more coherent set of clustered peptides.

### Removing peptides that are difficult to synthesize

The chemical nature of some amino acids, as well as the numerous steps and chemical compounds involved in synthesizing peptides, make some peptide sequences difficult to synthesize^49^. To overcome this, we excluded all of the peptides in our putative solution set that contained specific motifs from the superbinder pool as listed in **Table 2**.

### Validating putative superbinders using additional MHC predictors

To further filter our putative superbinder set, we utilized two additional MHC binding predictors to predict the binding affinity of each of the peptides in our set. Specifically, we used netMHCpan 4.0^26^ and MHCflurry^29^ to further filter our peptide list. For all three prediction methods, we computed the percentile rank of each peptide across all 12 alleles, and then computed the average percentile rank of the 8 alleles with the lowest values. Using this score, we selected 3000 top ranked peptides from each of the three predictors, and then chose the intersection of the peptides shared by all three predictors.

### Excluding self peptides

In a healthy state, HLA class I molecules present self-peptides sampled from intracellular proteins on the cell surface. These peptides are typically ignored by T-cells that have undergone negative selection — the process during T-cell development in the thymus that eliminates autoreactive T cells — thereby preventing immune responses against self^50,51^. To increase the probability that our superbinders will be immunogenic, we filtered out all putative superbinders that were similar to the self proteome. Specifically, we sliced all of the human peptidome into overlapping 9-mers and compared each of them to our putative superbinders using the Needleman-Wunsch global alignment algorithm. Peptides that shared an exact match of 7 or more amino acids with any self-peptide, as well as those with 6 identical amino acids and an additional 2 or 3 similar residues, were excluded from our putative superbinder set. The human peptidome used for this filtering was version 40, downloaded from https://www.gencodegenes.org.

### Generating a phylogenetic tree of the putative superbinders

We used the Molecular Evolutionary Genetics Analysis X (MEGA X) software package to further reduce our candidate peptide set from 991 to 100 peptides^32^. We created a phylogenetic tree of the peptides based on a maximum likelihood criteria, which estimates the likelihood of each tree by assigning probabilities to every possible evolutionary change at informative sites and identifying the topology that maximizes the overall likelihood. These methods incorporate models of amino acid or nucleotide substitution rates to guide the evaluation process^52^. The parameters used to construct the phylogenetic trees were as follows: the initial tree was generated using the Neighbor-Joining algorithm, and the amino acid substitution model applied was the JTT (Jones–Taylor– Thornton) model, with bootstrap analysis performed using 500 replicates. We then used hierarchical clustering using the bootstrap method.

### Runtime optimization

To improve the efficiency and scalability of the simulation we used multiprocessing. Multiprocessing allows computations to be distributed across multiple processors, which can work on different parts of a problem concurrently and takes advantage of modern hardware^53^. Specifically, the *superHLA* algorithm was run utilizing 8 cores, with simulations running in parallel, which significantly reduced the algorithm’s runtime. The algorithm was implemented in Python.

### HLA peptide binding assays

Biochemical measurement of peptide binding to class I MHC was performed using classical competition assays with purified MHC molecules and high affinity radiolabeled probe peptides, as detailed elsewhere^34^.

---

**Supplementary Figure 1.**
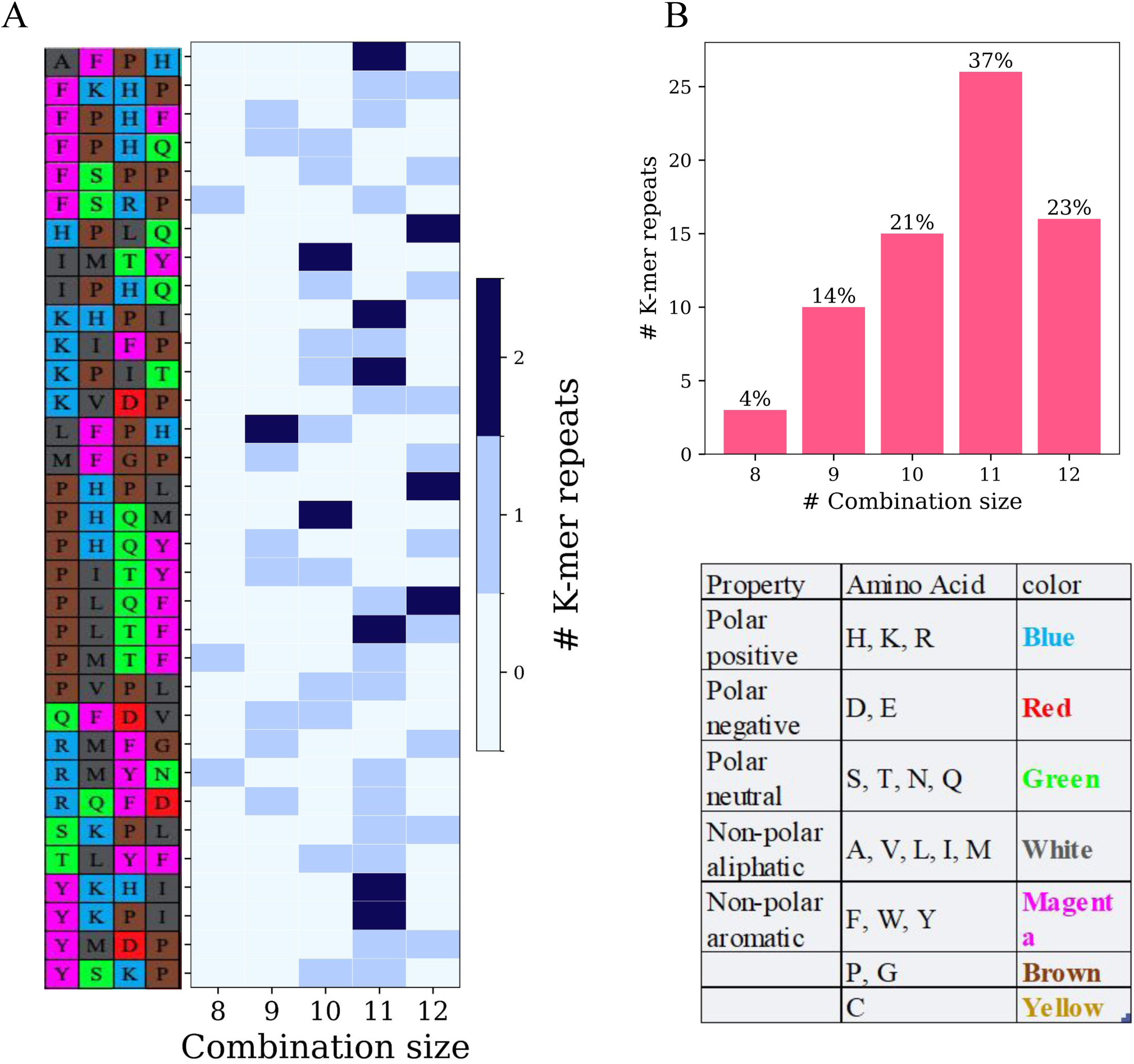
Frequent 4-mers in the superbinder validation set. **(A)** The set of 4-mer peptides that appeared in more than one peptide in the superbinder validation set. The biochemical properties of each amino acid are presented on the left. The x-axis represents the combination size of the final peptide pool.The number of 4-mer repeats within each combination is denoted by the color. (**B**) Percentage of the 4-mere repeats within different supertype combination sizes. The combination of 11 HLA representatives had the highest number of 4-mere repeats.

**Supplementary Figure 2.**
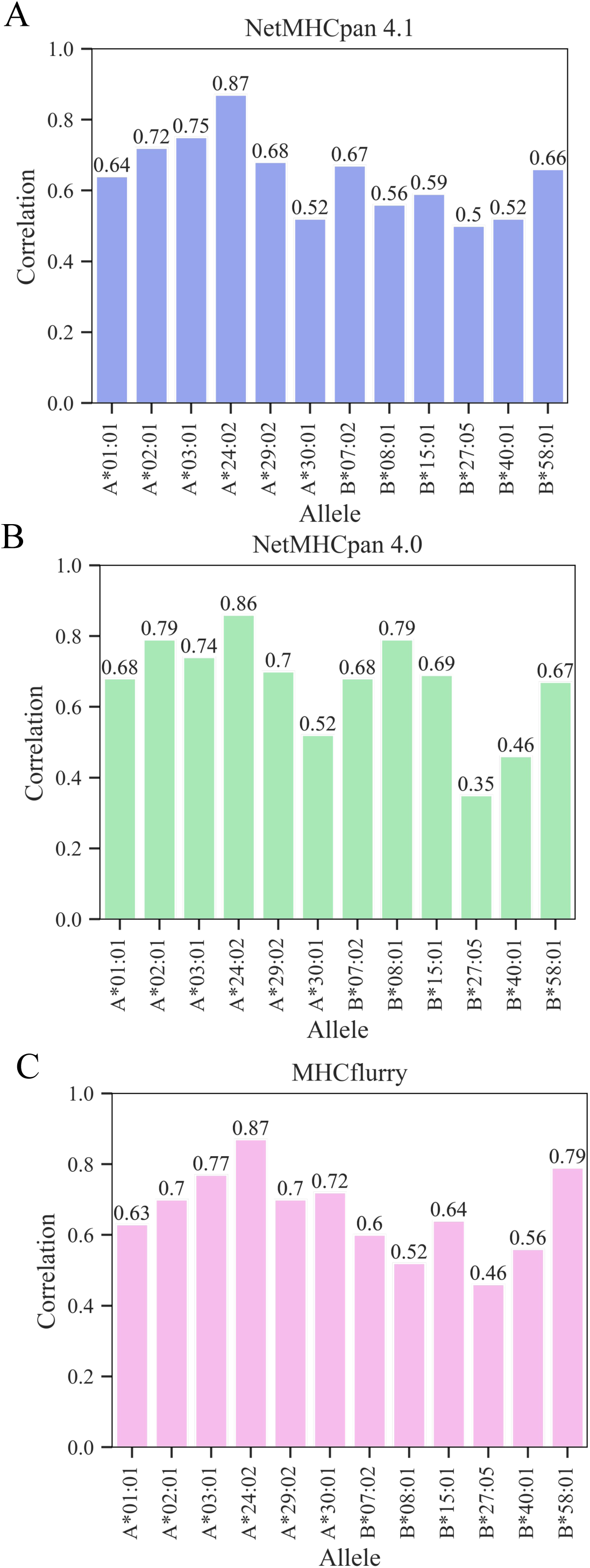
Distribution of Spearman correlation between the predicted percentile score to the experimental bonding assay values across the different HLA representatives within the different MHC1 predictors: (**A**) NETMHCpan 4.1 (**B**) NetMHCpan 4.0 and (**C**) MHCflurry.

**Supplementary Figure 3.**
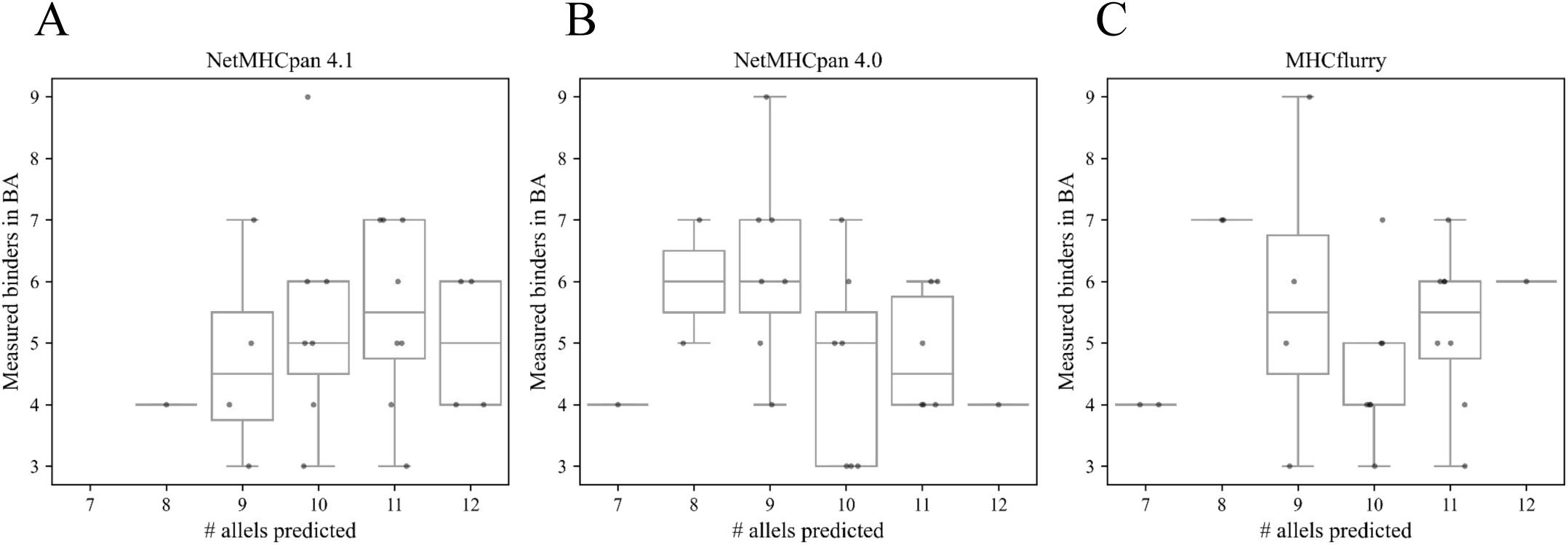
Number of the predicted HLA binders vs the number of HLA binders measured in experimental binding assays for each of the three predictors: (**A**) NETMHCpan 4.1 (**B**) NetMHCpan 4.0 and (**C**) MHCflurry.

**Supplementary Figure 4.**
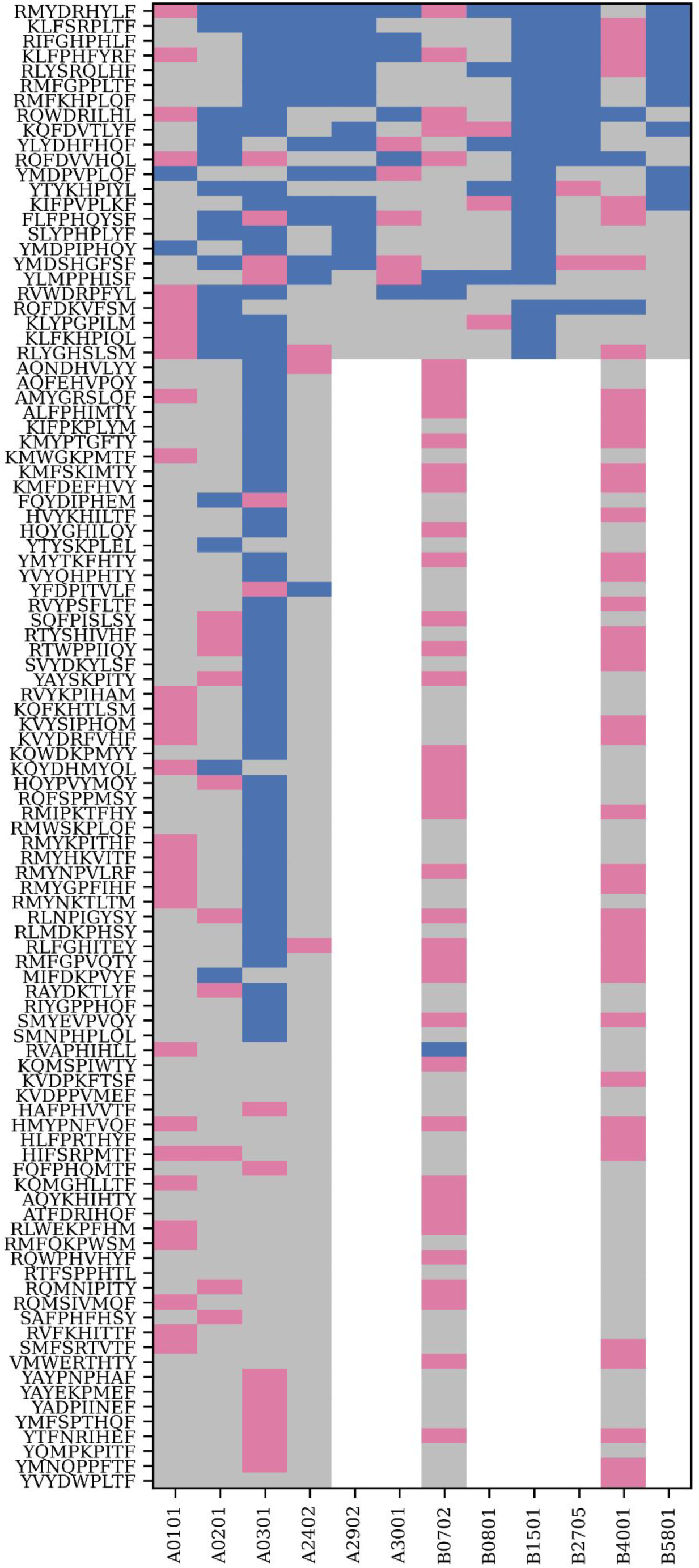
Classification heatmap of predicted vs. experimental peptide binding outcomes. Each cell represents the binding status of a peptide-allele pair, where colors indicate the agreement between in silico predictions and binding assay results: predicted binders that were validated (blue); predicted non-binders that were validated (pink); mismatched predictions and experimental measurements (gray) and combinations that were not tested experimentally (white) Rows are sorted by the number of consistent binding pairs (blue), highlighting consensus superbinders. Only 24 peptides were tested across all 12 HLA alleles across all supertypes.

